# Distinct muscle stem cell fates correlated with hyperplasia and hypertrophy during skeletal muscle growth in rainbow trout

**DOI:** 10.64898/2026.01.28.702282

**Authors:** Sabrina Jagot, Candice Babarit, Nathalie Sabin, Fabien Le Grand, Karl Rouger, Jean-Charles Gabillard

## Abstract

**Background:** In vertebrates, skeletal muscle grows postnatally through different strategies. While mammals predominantly rely on fiber hypertrophy after birth, many teleost fish retain the unique ability to generate new fibers via hyperplasia well into juvenile stages. The molecular mechanisms linked to the transition between hyperplastic-hypertrophic and hypertrophic muscle growth modes in fish remain poorly understood.

**Results:** We generated a single-cell transcriptomic atlas of muscle-derived cells from juvenile *Oncorhynchus mykiss* (rainbow trout) at five growth stages. Fifteen tissue resident cell populations were identified, including eight myogenic subpopulations spanning from quiescent stem cells to terminally differentiating myocytes. Two distinct transcriptional trajectories were uncovered thanks to RNA velocity analysis: one present only during hyperplastic growth and another maintained throughout growth, raising the possibility of specialization of muscle stem cells toward hyperplasia or hypertrophy. Comparative analyses with human single-cell atlases suggest that some trout myogenic subpopulations correlated with hyperplastic or hypertrophic muscle growth share similarities with human stage-specific (fetal or adult) myogenic subpopulations. Strikingly, we identified a population of *pax7*^+^/*pdgfrα*^+^ cells, indicating plasticity toward fibroblastic lineage and associating these cells with hypertrophic growth. Furthermore, both intrinsic changes in muscle stem cells and extrinsic remodeling of the extracellular matrix accompanied the skeletal muscle growth, suggesting a dynamic crosstalk between myogenic and mesenchymal compartments.

**Conclusions:** Our findings reveal the existence of two transcriptionally distinct muscle stem cell fates that correlate with hyperplastic and hypertrophic muscle growth in trout. The identification of a tissue-resident *pax7*^+^/*pdgfrα*^+^ subpopulation provides new insights into muscle stem cell plasticity and niche remodeling. This work establishes a comparative framework to explore the regulation of postnatal muscle growth across vertebrates.

## BACKGROUND

Skeletal muscle is mainly made up of bundles of individual fibers formed during embryonic and fetal development. Except during muscle regeneration following an injury, hyperplasia that corresponds to the formation of new muscle fibers, ceases around birth in mammals [1]. In contrast, a fascinating trait of most teleost is the persistence of hyperplastic muscle growth after hatching. Thus, a 200-fold increase in muscle fiber number is observed between the hatching embryo and a 4 kg salmon [2]. After an exponential muscle growth observed during the post-larval phase, hyperplasia sharply declines during the juvenile stage, underlining the existence of a very finely orchestrated regulatory mechanism over time [3]. Recently, a thorough analysis of the hyperplasia rate during the muscle growth of juvenile trout shows that the decline of this specific modality takes place earlier than previously thought, around 500 g in weight [4]. Thus, in salmonids, muscle growth occurs first through concomitant hyperplasia and hypertrophy modalities, then solely through hypertrophy. This highlights that the intensity and the duration of post-larval hyperplastic growth are crucial for the potential development of fish muscle mass. Both types of muscle growth modalities are ensured by the muscle stem cells (MuSC), also called satellite cells (SC), which are located between the basal lamina and the plasma membrane of the muscle fiber [5]. Once activated, MuSC proliferate and generate myogenic descendants that fuse either with existing muscle fibers to participate in the nuclear accretion required for fiber hypertrophy [6,7] or with each other to generate a new fiber and thus contribute to hyperplasia [8–11]. In mammals, it has been suggested that the end of ability for MuSC to form new fibers correlates with their positioning beneath the basal lamina [12]. However, in fish, establishment of the basal lamina and addressing of the MuSC to their niche occurs well before the end of hyperplastic growth [13]. Their behavior is therefore particularly fundamental in determining the balance between hyperplastic and hypertrophic muscle growth in fish.

In mammals, increasing evidence suggests the existence of subpopulations of MuSC with varying capacities for activation, proliferation, and self-renewal [14–16]. In elderly mice, impaired muscle regeneration capacities have been associated with changes in the properties of MuSC subpopulations [17]. During muscle regeneration, several subpopulations have been identified by single-cell RNA sequencing (scRNA-seq) analyses, which correspond to different states from quiescence to proliferation and differentiation [18,19]. In addition, distinct cell differentiation trajectories attributed to certain subpopulations were described, highlighting their specific fates. For example, during muscle regeneration, once activated, MuSC can adopt distinct trajectories, either returning to quiescent Pax7^+^ MuSC, committing to early activated Pax7^+^/Myf5^+^/MyoD1^-^MuSC, becoming proliferative Pax7^-^/Myf5^+^/MyoD1^+^ myoblasts or differentiating into fusion-competent myogenin (MyoG)^+^ late differentiated cells [18–20]. At the end of myogenesis, myocytes are differentiated mononucleated myogenic cells (expressing contractile genes such as myosin) able to fuse and form myofibers as multinucleated fibers. The term myogenic progenitor was chosen here to refer to cell that express *pax7* from quiescent to early activated MuSC, and the one myogenic precursor to cell involved in myogenesis from *pax7* expression to *myogenin* expression.

Furthermore, the myogenic capacities of MuSC are regulated by the interplay between intrinsic (cell-autonomous) and extrinsic signals originating from their niche (non-cell-autonomous) [21]. In this respect, in addition to muscle fibers, the fibro-adipogenic progenitors (FAP) and the macrophages have been clearly identified as cellular players capable of influencing the fate of MuSC [22–24]. Given this level of complexity in the regulation of myogenesis, numerous studies in mammals have recently focused on the characterization of all the muscle-derived cells (MDC) [25] and their potential interactions in various contexts, such as disease [26,27], regeneration [28,29], and aging [17,30]. However, no equivalent approach has yet been conducted to describe the decline in hyperplasia observed during muscle growth in fish with a focus on MDC at single cell resolution.

In the present study, we aimed to characterize MDC dynamics during trout muscle growth in order to identify the mechanisms underlying the transition from combined hyperplasia and hypertrophy-based to only hypertrophy-based muscle growth. In particular, we sought to determine whether and how many subpopulations of myogenic cells co-exist in the tissue and, if so, how they evolved during muscle growth. For that purpose, we performed scRNA-seq analysis on MDC extracted at five growth stages between 10 g and 1.5 kg, exhibiting progressively decreasing levels of hyperplasia [3]. This has enabled us to generate the first atlas of MDC in fish, providing a wealth of data for in-depth analysis of myogenic and mesenchymal cells. From these original findings, we identified eight myogenic cell subpopulations corresponding to distinct states of commitment. Importantly, analysis of their transcriptomic trajectories allows us to identify two myogenic trajectories correlated with either hyperplastic or hypertrophic muscle growth. For the first time, we present an original two-branch myogenesis pattern in fish and the presence of a tissue-resident *pax7*^+^/*pdgfrα*^+^ population.

## METHODS

### Animals

Rainbow trout (*Oncorhynchus mykiss*) were maintained in a recirculating system under a simulated natural photoperiod at 12 ± 1 °C (pH 7.8–8.4; NH₄ < 0.1 mg/L). Fish were fed daily to apparent satiation with a commercial diet (Le Gouessant). All animals were reared at the experimental facilities of the INRAE Fish Physiology and Genomics Laboratory (LPGP, Rennes, France; https://doi.org/10.15454/45d2-bn67; authorization number D35-238-6), accredited by the French veterinary authorities. All animals used in the present study were juvenile and didn’t undergo sexual maturation. For tissue sampling, fish were anesthetized with 50 mg/L tricaine methanesulfonate (MS-222) and subsequently euthanized with an overdose of 200 mg/L tricaine. All experimental procedures were conducted in strict compliance with the European Directive 2010/63/EU on the protection of animals used for scientific purposes. The euthanasia protocol was approved by the Ethical Committee for Animal Experimentation of Rennes (CREEA).

### Cell suspension collection for scRNA-seq experiment

Mononucleated cells were isolated from white muscle juvenile trout (10 g to 1.5 kg body weight) as previously described [31]. Information related to sample collections as well as correspondence between weight, age and length of fish were provided (Table S1). Briefly, 20 to 80 g of white muscle were mechanically dissociated with scalpels and enzymatically digested by collagenase (Sigma, #C9891) and trypsin (Sigma, #T4799) prior to filtration (Falcon Cell Strainer 100 μm, #2360 and 40 μm #2340). In order to remove as much debris and red blood cells as possible from cell suspension, a 20%-60% Percoll gradient was applied. Cells were resuspended in Dulbecco’s modified Eagle’s medium (DMEM) containing 10% fetal calf serum (FCS; Sigma, #F7524) and 1% antibiotic–antimycotic solution (Sigma, #A5955) and conserved at 4°C, overnight (ON). Cells were washed 3 or 4 times with DMEM plus 1% of antibiotics and filtered using 40 µm Flowmi® Cell Strainers. After ensuring a cell viability greater than 70% and a concentration at 700 to 800 cells per µL, cell suspensions (∼12,000 cells per reaction) were loaded onto the Chromium system (10X Genomics). Following the instructions of the manufacturer, cDNA amplification and scRNA-seq libraries were constructed using Single-Cell 3′ Gene expression kit v3.1 (10X Genomics). Twelve to 14 PCR cycles were used to amplify cDNA libraries and quantified on a Qubit Fluorometer (Thermo Fisher Scientific). Average fragment size was determined on a Bioanalyzer (Agilent 2100). The libraries were sequenced using the Illumina Novaseq 6000 on a flowcell S4 200 cycles v1.5 (28 cycles for Read 1, 150 cycles for Read 2, 10 cycles for i7 index and 10 cycles for i5 index).

### Single-Cell RNA-seq data processing

Reads obtained from scRNA-seq were mapped on the *Oncorhynchus mykiss* reference genome (USDA_OmykA_1.1; NCBI ReSeq Assembly: GCF_013265735.2). Table of Official NCBI Gene Names, Gene Description and gene names used in this article are available in Table S2. The sequencing depth was 157,777 reads per cell and count matrices were generated using CellRanger v7.0.1 (10X Genomics). The obtained count matrices were analyzed using Seurat v.4.3.0. Classical quality filters were used, *i.e*., the droplet containing cell have been kept i) if the number of gene expressed in droplet was higher to 500 and less than 10,000 ii) if the proportion of mitochondrial genes was not exceeded 6% and iii) if the proportion of ribosomal associated genes was higher than 2%. Droplets containing more than one cell were identified and remove from the count matrices using DoubletFinder [32]. The raw UMI count were normalized using LogNormalize method, the top 4,000 highly variables genes across all cells for MDC atlas dataset (top 3,000 highly variables genes across all cells for myogenic or mesenchymal datasets) and the 30th first principal components were used for building the shared nearest-neighbor graph and clustering were done with Louvain method (resolution: 0.15, 0.3 and 0.4 for MDC atlas, myogenic subset and mesenchymal subset, respectively). Datasets from different samples were integrated and batch effects were corrected using RunFastMNN function from SeuratWrappers package (v.0.3.1). Cell populations were visualized using uniform manifold approximation and projection (UMAP) algorithm implemented in Seurat package. For all clusters, differential analyses were made to find marker genes using FindAllMarker function (MAST test: logFC ≥ 0.25; min.pct = 0.25; min.diff.pct = 0.2; Bonferroni adjusted p < 0.01). Top 100 gene markers for each cluster of myogenic subset and mesenchymal subset were used to annotate cell type/states (Tables S3 and S4). As DESeq2 [33] Likelihood Ratio Test fits well with longitudinal or temporal studies, to identify differentially expressed genes that shared patterns across weight conditions, we used DESeq2 R package (test: LRT with adjusted p-value < 0.05, v.1.38.3). We grouped these differentially expressed genes into distinct expression profiles, using DEGreport package (v.1.34.0), only groups containing 15 genes at least were kept. We focused on genes showing a relatively linear deregulation with weight: groups 3 and 4 in myogenic cell cluster 0 (1146 DEG), groups 1 and 5 in myogenic cell cluster 1 (461 DEG), groups 3 and 4 in mesenchymal cell cluster 1 (881 DEG) and groups 2 and 4 in mesenchymal cell cluster 3 (675 DEG). To visualize specific gene expressions, we applied the imputation ALRA method [34]. All of these analyses were performed using R software (v4.2.3).

### Statistical analysis

Differential cell-type proportion analysis was performed using the scanpro framework [35], which implements a compositional modeling strategy tailored for single-cell datasets with biological replication. For each sample, cluster proportions were computed from the total number of cells per replicate. A linear model analogous to ANOVA was fitted to each cluster to test for differences across weight groups, with weight treated as a categorical variable. This approach ensures that biological replicates, rather than individual cells, constitute independent observations. P-values were corrected for multiple comparisons using the Benjamini–Hochberg method.

### Velocity analysis

The command line tool velocyto (v.0.17) was used to generate loom files containing spliced and unspliced matrices from CellRanger outputs. Implemented on Python (v3.9), scvelo (0.2.5) and scanpy (1.9.3) modules were used to read and concatenate loom files and then, merge with the anndata object obtained after Seurat analysis described above. Temporally unified RNA velocity for single-cell trajectory inference (UniTVelo) [36], implemented on Python (v.3.7) and TensorFlow (v.2.10, for GPU using), was used for transcriptomic trajectory analysis and identification of genes that could drive cell fates. The model estimates velocity of each gene and updates cell time based on phase portraits concurrently. Default parameters were used to construct the UniTVelo model for myogenic subsets of 10 g, 100 g or 1000 g samples. To construct the UnitVelo model for myogenic subset with all sample, adjusted parameters were used (NUM_REP=2; NUM_REP_TIME=’re_init’; N_NEIGHBORS=50; N_TOP_GENES=3000).

### Homo sapiens and Oncorhynchus mykiss scRNA-seq comparisons

Human [37,38,39] and Trout datasets were compared using the SAMap [40] (v.2.0.0) algorithm, a method for mapping scRNA-seq datasets between distant species implemented on Python (v.3.9). First, reciprocal Blast searches between the two transcriptomes of the two species were realized to construct a gene–gene bipartite graph with cross-species edges that connecting homologous gene pairs. Second, the graph was used to project the two datasets into a UMAP. Iteratively, the expression correlation of homologous genes was used to update the bipartite graph and relax SAMap’s initial dependence on sequence similarity. A mapping score (ranging from 0 to 1) was computed among all possible cross-species cluster pairs (Table S5 and S6 to S9 for MDC atlas and myogenic subset comparisons, respectively). For myogenic subset comparisons, human datasets of myogenic cell subsets from adult and developmental stages were concatenated before SAMap analysis with all myogenic subset cells from the 12 scRNA-seq samples of trout white muscles.

### RNAscope experiments

To visualize the spatial distribution of mRNA of interest in trout muscle, *in situ* hybridization was performed using the RNAScope V2 kit (Bio-Techne, #323100) according to the manufacturer’s protocol for formalin-fixed paraffin-embedded (FFPE) sections. In brief, white muscle samples were fixed with 4% paraformaldehyde (ThermoScientific, #28906) (4°C, ON) and embedded in paraffin. FFPE cross-sections were cut at 4 µm, baked (60°C, 1 hour), deparaffinized and air dried. After 10 min in hydrogen peroxide solution (Bio-Techne, #322335), sections were successively treated with 1X Target Retrieval (Bio-Techne, #322000) (100 °C, 15 min) and Protease Plus solution (Bio-Techne, #322331) (40 °C, 18 min). Then, sections were hybridized with the respective RNAScope probes (40 °C, 2 hours). All probes used in this study were provided by Bio-Techne. They were *pax7* (Om-pax7b-cust, #575461) which stained for transcripts from all *pax7* paralogs genes [13], *myod1* (Om-myod1-C2, #1039251-C2), *myod2a* (Om-myod2-C3, #1039261-C3), *pdgfrα* (Om-pdgfra-C2, #1029301-C2) and *pdgfrβ* (Om-LOC110488323-C2, #1567191-C2). All probes were designed with the cDNA FASTA sequences from *pax7b-LOC100884157, myod-LOC100136773*, *myod2-LOC100136782*, *pdgfra-LOC110521671* and *LOC110504831*. The DapB (4-hydroxy-tetrahydrodipicolinate reductase) was used as the negative probe (#320871). Negative probe staining and enlarged examples of staining were illustrated in Figure S1 and Figure S2, S3, respectively. The sections were stored in 5X SSC buffer (Invitrogen, #15557044) ON, and the amplification and multiplexing steps took place the next day. Multiplexing was performed using Opal multiplex detection system (Akoya Bioscience, Opal 520 #FP1487001KT, Opal 570 #FP1488001KT, Opal 690 #FP1497001KT). Upon completion of the amplification and multiplexing steps, the sections were incubated with DAPI and cover-slipped with Mowiol (Millipore, #475904). Sections were imaged within one week of staining using Axio Scan z1 slide scanner (Zeiss).

## RESULTS

### Trout muscle-derived cell atlas is distinct from that of human cells by specific enrichment of myoblasts during hyperplastic-hypertrophic growth

To shed light on the nature of the cell involved in growing juvenile trout muscle, a scRNA-seq analysis was performed on MDC freshly isolated from the white muscle of trout weighing 10 g, 100 g, 500 g, 1 kg and 1.5 k g (Figure 1a). These stages encompass the transition from hyperplastic/hypertrophic growth to hypertrophic growth [4]. After quality filtering, 34,277 cells with an average of 3,001 expressed genes per cell were retained, constituting the first atlas of MDC in fish. The uniform manifold approximation and projection (UMAP) [41] revealed the presence of 15 cell populations (Figure 1b), which were identified by specific markers for each cluster, using MAST tests (Figure 1c). Three were associated with skeletal muscle, with *itga7* and *pax7b* as marker genes of MuSC, *myog* identified a myoblast cluster and *myl1b* and *tnnc2* fast myocytes [28]. Two cell populations were related to connective tissues (mesenchymal cells and tenocytes) with *pdgfrα* [42] and *tnmd* [43] as marker genes. Two clusters originated from vascular tissues (endothelial cells and smooth myocytes), six corresponded to blood cells (platelets, macrophages, T cells, B cells, granulocytes, and erythrocytes), and one to Schwann cells. More than half of MDC are represented by myogenic cells (29%; represented by MuSC, myoblasts and fast myocytes clusters) and mesenchymal cells (26%). Tenocytes and Schwann cells are present in MDC at 13% and 9%, respectively. To identify any specificities of trout MDC diversity, the atlas generated was compared with that already established with cells derived from diaphragm muscle of adult human [37] by self-assembling manifold mapping (SAMap), a method dedicated to detect homologous cell types [40]. The UMAP projection of the inter-species manifold (Figure 1d) and the distribution of mapping scores between the two atlases (Figure S4a-d and Table S5) showed a high degree of correspondence with a global mapping score of 0.67. Fast myocytes represented 5% of trout MDC but 1% of human ones, as a reflect of differing muscle types in both organisms. Interestingly, the proportion (16%) of MuSC in the trout atlas was identical to that of skeletal muscle satellite stem cells in the human atlas. One major difference between the two MDC atlases was the presence of 8% of myoblasts engaged in myogenic differentiation in the trout, that was absent in human. The distribution of scRNA-seq samples in hierarchical clustering was performed on the averaged expression profiles of each sample. The resulting dendrogram clearly identified two main blocks (Figure 1e): one between 10 g and 100 g and one from 500 g to 1.5 kg. Analysis of cell population proportions (Figure 1f and S4e) showed that MuSC remains relatively stable (15% ± 3%, Figure S4f) throughout the entire period, whereas myoblasts decrease from 16% to 3% between 10 g and 1.5 kg (Figure 1f and S4g). At the same time, proportion of endothelial cells significantly increases (2% to 18%, Figure S4h). These findings show that muscle growth is accompanied by a progressive remodeling of the relative cell population proportions.

**Figure 1.**
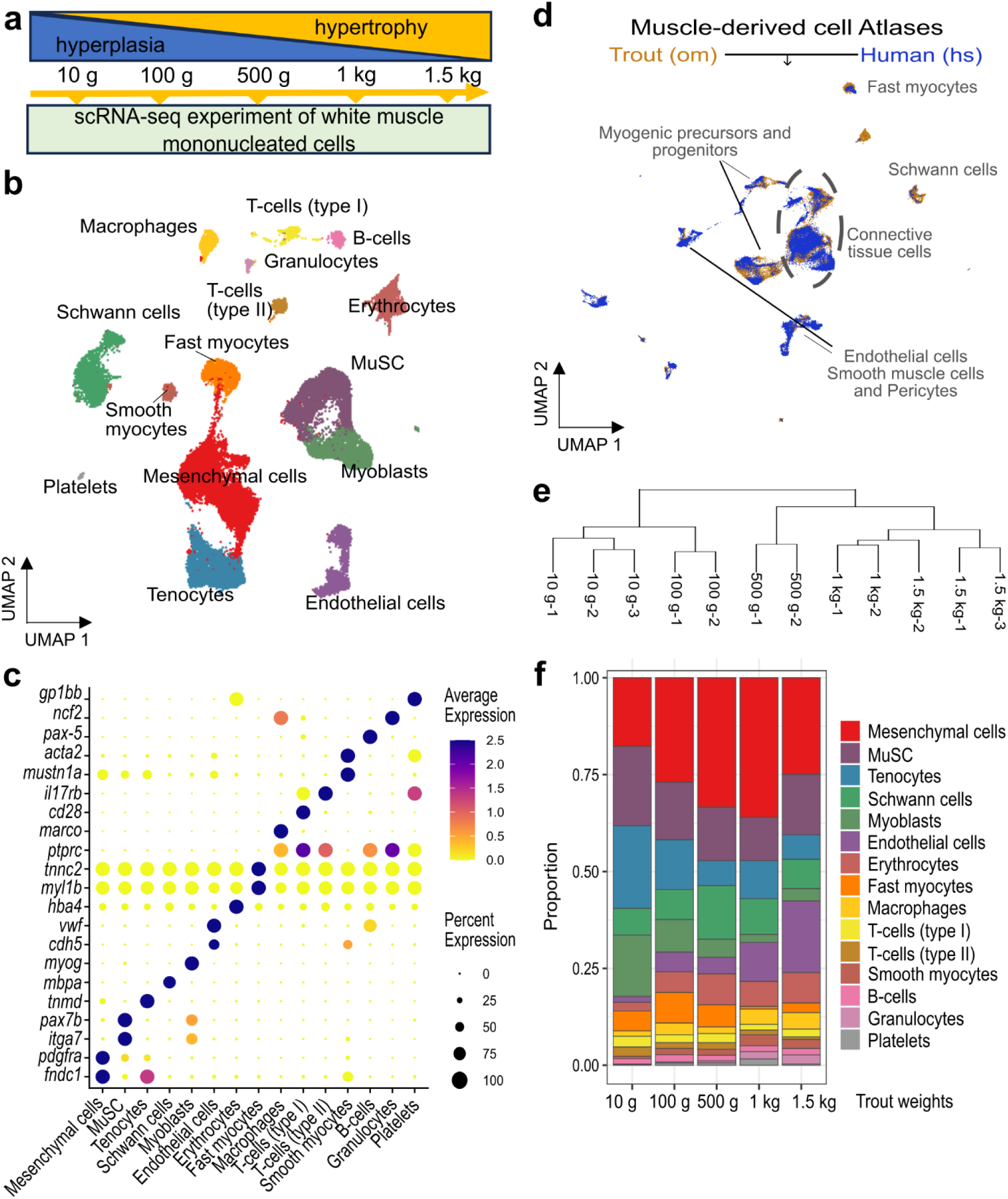
Characterization of muscle-derived cell diversity in juvenile trout muscle during growth. (a) Overview of the single-cell RNA sequencing (scRNA-seq) experiment done on muscle-derived cells (MDC) isolated from white muscle of trout at different growth times (between 10 g and 1.5 kg). (b) Uniform manifold approximation and projection (UMAP) showing the clustering of 34,277 cells into 15 distinct cell populations (all samples and all stages merged). Each cluster is color-coded and annotated based on expression of marker genes. (c) Dot plot of average expression of key marker genes in each cluster. Size and color of the dot reflect the percentage of cells in the cluster and the average expression, respectively. ALRA corrected matrix used for expression levels representations (d) UMAP showing the cross-species comparison between atlas of trout (om) and human [35] (hs) MDC, using SAMap [37]. (e) Pair-wise correlation of averaged gene expression profiles of scRNA-seq samples visualized as dendrogram. (f) Evolution of the proportion of MDC populations during growth.

### Trout myogenic program is originally divided into two distinct transcriptional trajectories

To further characterize myogenic cells, we performed clustering combined with MAST test-based differential analysis, which enabled the identification of eight subpopulations (clusters 0 to 7) (Figure 2a, b, Table S3). Clusters 0 and 1 expressed *itga7* and the MuSC marker *pax7b* [44,45]. They corresponded to the undifferentiated myogenic cells, and they would later be called progenitors. The expression of *myf5a* and *myod2a* attributed a characteristic signature for cluster 1, suggesting a distinct early activated state of these myogenic progenitor cells. Interestingly, the *myod* paralogs showed very singular expression profiles depending on the cluster. Thus, *myod1* was not expressed in clusters 0 and 1, but strongly in cluster 2, and moderately in clusters 4 and 5. *myod2a* paralog was strongly expressed in cluster 1 and detected in clusters 4 and 5, which also showed high levels of expression for *myod2b* (with *myod2b* as marker of cluster 5 cells). Finally, it should be noted that the *myod* paralogs were not expressed by clusters 3, 6 and 7. The genes encoding the late differentiation marker *myog* and *myomaker* (*mymk*) were also expressed by cells in the cluster 2, corresponding to fusion competent myoblasts [46]. Cluster 3 and 6, was defined by the expression of the *myosin light chain 1* gene (*myl1*), indicating that it is composed of myocytes. Cluster 4 was characterized by the expression of *mki67* and cell cycle genes, suggesting its enrichment in highly proliferative cells. Finally, cluster 7 was mostly composed of genes encoding muscle myosins and markers of slow muscle. Following this mapping of myogenic cell states, we sought to determine whether the distinct cell fates identified were dynamically associated. For that, RNA velocity analyses have been conducted with UniTVelo tool [36], which predicts transcriptional dynamics and infer future cell states by measuring newly transcribed pre-mRNAs (unspliced) and mature mRNAs (spliced). Over the entire dataset, the transcriptional trajectories first showed a 3-level stratification with a point of origin corresponding to the myogenic progenitors of clusters 0 and 1, an intermediate state composed of the early committed myogenic cells of clusters 4 and 5 and a final state with the late differentiating cells of cluster 2 (Figure 2c). This cell segmentation was illustrated by the “latent time”, representing the cell’s internal clock and its position in an underlying biological process, where each cell was positioning from root to ending (Figure 2d). Indeed, complementary to the first representation of the model generated by UniTVelo, the myogenic progenitors were positioned in clusters 0 and 1, while the cells downstream of this process were the cells in cluster 2, corresponding to those at the end of myogenic differentiation. Cluster 5 cells were uniformly positioned in an intermediate position in this myogenic differentiation process. Interestingly, the cells in cluster 4 appeared to be composite, with cells clearly identified as intermediates to the right of the UMAP and cells positioned more at the end of the process. These cells appeared close to cluster 2 both spatially on the UMAP and on latent time. Thus, the RNA velocity analysis identified two distinct trajectories of myogenic differentiation, from 0/1 cluster to cluster 2 via either cluster 5 or cluster 4.

**Figure 2.**
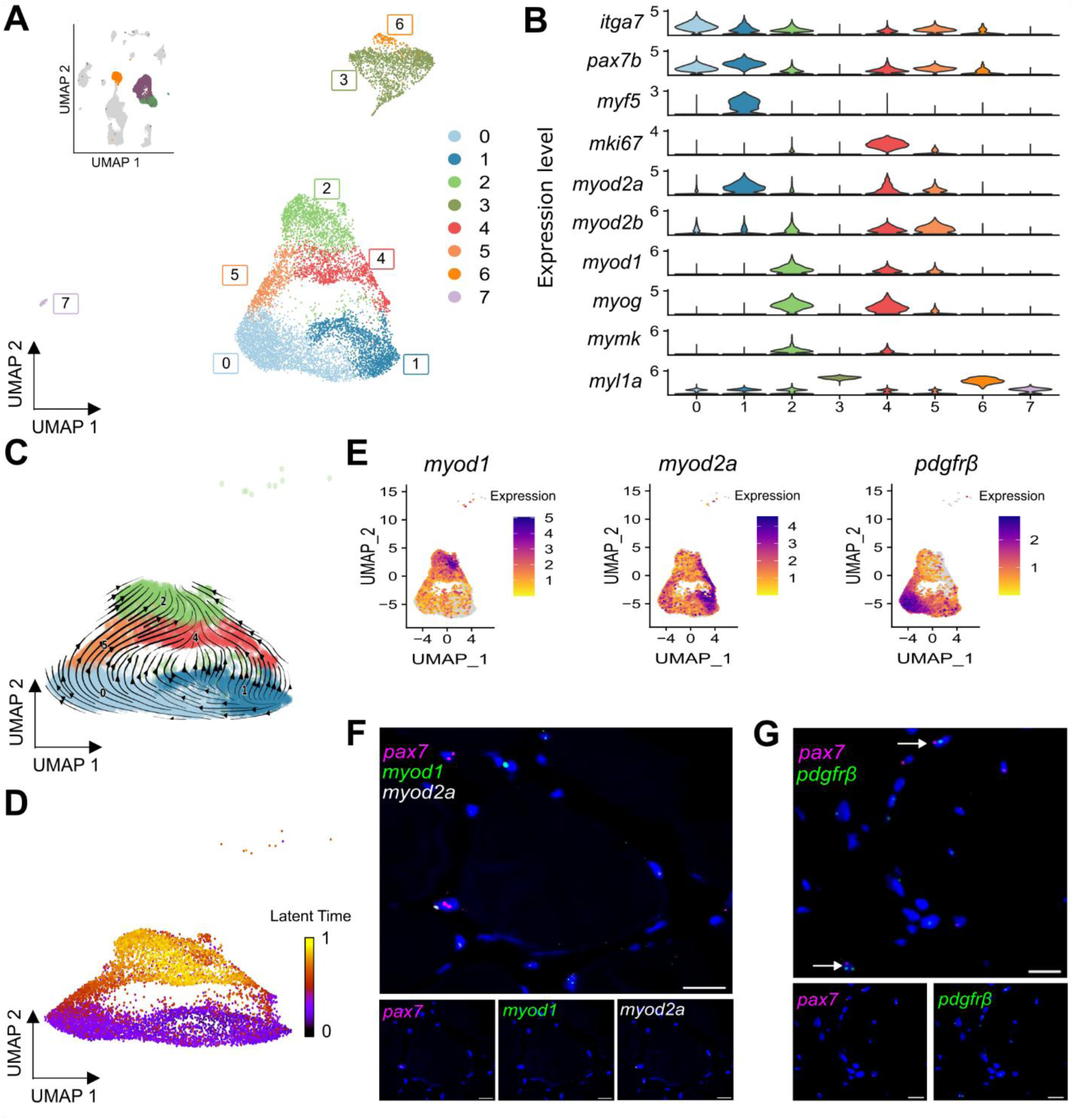
Identification of two transcriptional trajectories of myogenic differentiation during muscle growth in trout. (a) UMAP of myogenic cell subset (including muscle stem cells - MuSC, myoblasts and fast myocytes clusters, respectively showed in the top left inset in purple, green and orange) and a clustering identifying 8 distinct subpopulations. (all samples and all stages merged; 9,955 cells) (b) Violin plot of key marker gene expression profiles in clustered myogenic cells. (c) Transcriptional trajectories obtained from RNA velocities analysis of single-cell RNA sequencing (scRNA-seq) of myogenic precursors and progenitors (cluster 0, 1, 2, 4 and 5; all samples merged) depicted as streamlines. Clusters are shown in different colors. (d) Latent time projection of myogenic precursors and progenitors (cluster 0, 1, 2, 4 and 5; all samples merged). (e) Feature plot of *myod1*, *myod2a* and *platelet-derived growth factor receptor beta* (*pdgfrβ*). (f) RNAscope validation of distinct driver gene expression in white muscle cross-sections of 100 g trout. Representative images showing the expression of *pax7* (magenta), *myod1* (green) and *myod2a* (white). DAPI (blue) stains the nuclei. Scale bar = 20 µm. (g) RNAscope validation of *pdgfrβ* driver gene expression in white muscle cross-sections of 500 g trout. Representative images showing the expression of *pax7* (magenta) and *pdgfrβ* (green). DAPI (blue) stains the nuclei. Scale bar = 10 µm. The ALRA corrected matrix was used to visualize expression levels.

We then investigated which genes underlie these trajectories and mark the different states of myogenic cells *in vivo*. UniTVelo classified driver genes into three temporal categories: induced, repressed, or transient, differing by the timing of their peak expression according to latent time. These gene expressions were visualized with heatmaps (Figure S5 and Table S10). Among the induced genes (Figure S5a), we found genes associated with the myogenic differentiation process (such as *mymk*, *mef2cb*, and *myod1*). Phase portraits confirmed that *myod1* expression was positively correlated with latent time for both spliced and unspliced transcripts, with a more marked increase in unspliced transcripts (Figure 2e, Figure S5b). This is consistent with its higher expression in fusion-competent myoblasts (cluster 2). Overexpressed in cluster 4 cells, we found cell cycle genes (such as *cdk1*, *cdk2*, *mki67*, *cyclin D1* and *D2*, *cdc2 kinase*, and the *PCNA-associated factor*) among these induced genes. Repression driver genes, expressed early in the latent time, included *itga7*, *pax3-like*, *pax7a2*, and *col5a3a* (Figure S5c and Table S10). Surprisingly, *myod2a*, one of the homologous genes of *myod*, was among these early-expressed genes, overexpressed principally in cluster 1 cells and subsequently in clusters 4 and 5 (Figure 2e). Phase portraits also showed that *myod2a*, unlike its paralog *myod1*, was expressed inversely correlated with latent time (Figure S5d). To verify potential differential expression of *myod* paralogs in the myogenic cells, we performed RNAscope experiment for *pax7*, *myod1*, and *myod2a* on muscle tissue sections from 100 g trout (Figure 2f, Figure S2). We detected *pax7*^+^/*myod1*^-^/*myod2a*^-^, *pax7*^+^/*myod1*^+^/*myod2a*^-^, *pax7*^+^/*myod1*^+^/*myod2a*^+^, *pax7*^+^/*myod1*^-^/*myod2a*^+^, and *pax7*^-^/*myod1*^+^/*myod2a*^+^ cells, confirming *in vivo* the differential expression of *myod* paralogs by trout myogenic cells. The UniTVelo modeling identified 155 transient driver genes (Figure S5e and Table S10), among them *pdgfrβ*, which caught our attention as a marker of a recently identified subset of MuSC [47]. This gene was expressed predominantly in cells from clusters 0 and 5, for both spliced and unspliced transcripts (Figures 2e and S5f), making it a potential marker of the myogenic differentiation axis comprising cells from clusters 0 and 5. RNAscope confirmed the presence of *pax7*^+^/*pdgfrβ*^+^ myogenic cells in 500 g trout muscle tissue section (Figures 2g and S3). The *in vivo* validation of key gene expressions associated with these differentiation trajectories supported the idea of a cell specialization process specific to the hyperplastic growth of teleosts.

### Two myogenic transcriptional trajectories correlate with hyperplasia-hypertrophy or hypertrophy growth modality

To determine how the two transcriptional trajectories identified by RNA velocity evolved during muscle growth, we quantified the proportions of the 8 myogenic clusters across biological replicates at each weight stage (Figures 3a and S6a). Cluster 0 significantly increased with body weight (adjusted p = 0.005; Figure S6b). It represented approximately 27% of myogenic cells at 10 g, 100 g and 500 g, and rose sharply to 51% (±3%) at 1 kg and 47% (±10%) at 1.5 kg. Similarly, cluster 1 also increased significantly across weight stages (adjusted p = 0.007; Figure S6c), from 13% (±1%) at 10 g to 23±5 and 25±2% at 500 g and 1.5 kg, respectively. In contrast, cluster 4 showed a significant decrease with increasing weight (adjusted p = 0.005; Figure S6d). This intermediate state accounted for 16±8% of myogenic cells at 10 g but declined to 3±1% at 1 and 1.5 kg. Cluster 2 also displayed a significant reduction from 21±5% at 10 g to 11±4% at 1.5 kg. By comparison, cluster 5 did not show a statistically significant change across weight stages (adjusted p = 0.182; Figure S6e). Its proportion fluctuated between 7±2% at early stages and 5±1% at later stages. These quantitative analyses indicated a significant expansion of progenitors from clusters 0 and 1, a marked decline of cluster 4, and a reduction of cluster 2, while cluster 5 remains proportionally maintained. Stage-specific RNA velocity analyses further revealed remodeling of differentiation trajectories (Figure 3b-f). At 10 g and 100 g, two distinct differentiation routes were visible: one passing through cluster 4 and another through cluster 5. On 100 g trout, trajectories between clusters 0 and 1 emerged, showing the maintenance of the progenitor pool at this stage. At 500 g, the trajectory involving cluster 4 appeared reduced. At 1 kg and 1.5 kg, streamlines predominantly followed the cluster 5 route, whereas the branch through cluster 4 was strongly attenuated. Thus, the decline in cluster 4 abundance coincides with progressive loss of the corresponding transcriptional trajectory.

**Figure 3.**
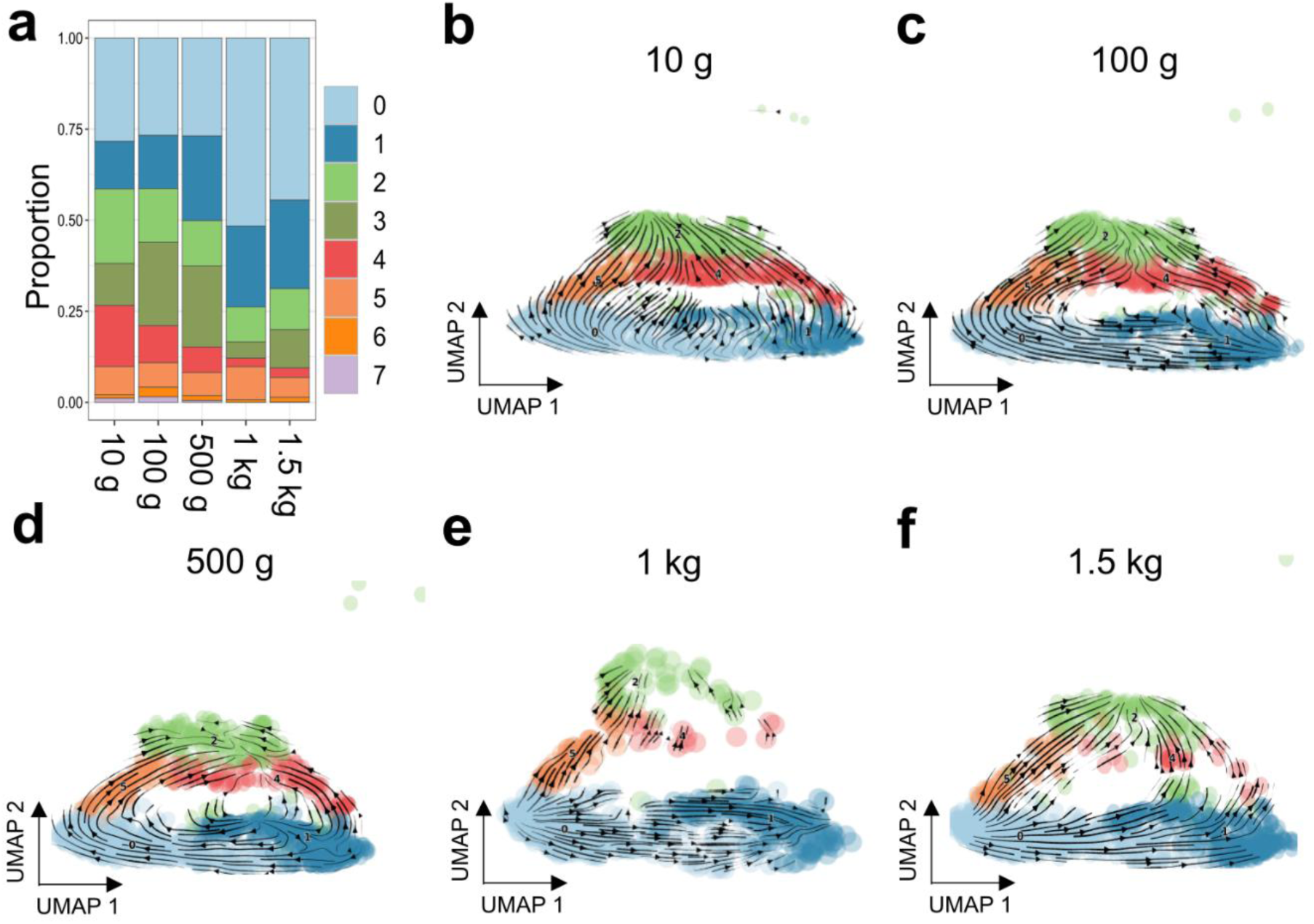
Evolution of the transcriptional trajectories during muscle growth in trout. (a) Evolution of the proportion of myogenic cell subpopulations during growth. (b-d) RNA velocities based on single-cell RNA sequencing (scRNA-seq) of myogenic precursors and progenitors (cluster 0, 1, 2, 4 and 5) depicted as streamlines for 10 g (b), 100 g (c), 500 g (d), 1 kg (e) and 1.5 kg (f) merged samples (n = 2-3). Clusters are shown in different colors.

To further explain this modeling of the myogenic differentiation dynamics in juvenile trout muscle growth, we carried out four detailed comparative analyses [40] between trout juvenile myogenic clusters with adult [37] and stage-defined embryonic [39] or fetal [38] human skeletal muscle datasets (Figures 4 and S7, Tables S6-S9). Cluster 4 consistently aligned with myogenic progenitor populations across developmental stages. Upon the comparison with the first wave of myogenesis (5.1–6.5 weeks), it showed highest mapping scores toward embryonic progenitors (hs_PAX3^+^PAX7^+^MyoProg, 0.36; hs_PAX3^+^MyoProg, 0.32; hs_PAX7^+^SPON2^+^MyoProg, 0.29). This embryonic progenitor similarity persisted upon the comparison with the beginning of the second wave (7.2–9 weeks), with moderate alignment to hs_PAX3+PAX7+MyoProg (0.29), hs_PAX7+NTN5+MyoProg (0.25) and to embryonic precursors hs_MyoB2 (0.24). When homology tests were done later at fetal stages (12–14 and 17–18 weeks), cluster 4 distributed between adult satellite cells (hs_14, ∼0.34) and fetal progenitors (hs_MP, ∼0.17–0.27; hs_SkM.Mesen, 0.21). In contrast, cluster 5 showed more variable alignment across development stage with embryonic progenitors during early stages (0.27 in the first wave; 0.26 in the second wave) and showing minor correspondence with adult satellite cells (hs_14, ∼0.14–0.16). When homology tests were done later at fetal stages, cluster 5 displayed a strong alignment with the adult satellite cell compartment (hs_14, 0.51), which remained stable across later time points. Across all analyses, differentiated trout cluster om_3 reproducibly aligned with human differentiated muscle populations (hs_7 and hs_MYL3+MyoC, ∼0.50–0.53), while trout progenitor cells from clusters 0 and 1 predominantly mapped to adult satellite cells (hs_14, ∼0.41–0.49), with a secondary similarity to fetal progenitors depending on stage for cluster 1. Taken together, these stage-resolved SAMap analyses demonstrate that the two trout intermediate states underlying the distinct RNA velocity trajectories exhibit differential cross-species alignment: when compared with datasets from adults and three different stages of the second myogenesis, cluster 1 and cluster 4 consistently shows similarity to embryonic and fetal myogenic progenitor populations, whereas cluster 0 and cluster 5 repeatedly aligns more strongly with adult satellite cell-associated compartments. Consistent with these cross-species positioning results, the quantitative remodeling of trout myogenic clusters across growth stages (Figures 3 and S6) reveals a marked decline of cluster 4, whereas cluster 5 remains proportionally maintained across stages. Thus, the intermediate state that shows stronger alignment with embryonic and fetal progenitor populations (om_4) is the one that significantly declines during muscle growth, while the state exhibiting stronger correspondence to adult satellite cell-associated compartments (om_5) persists throughout the growth period.

**Figure 4.**
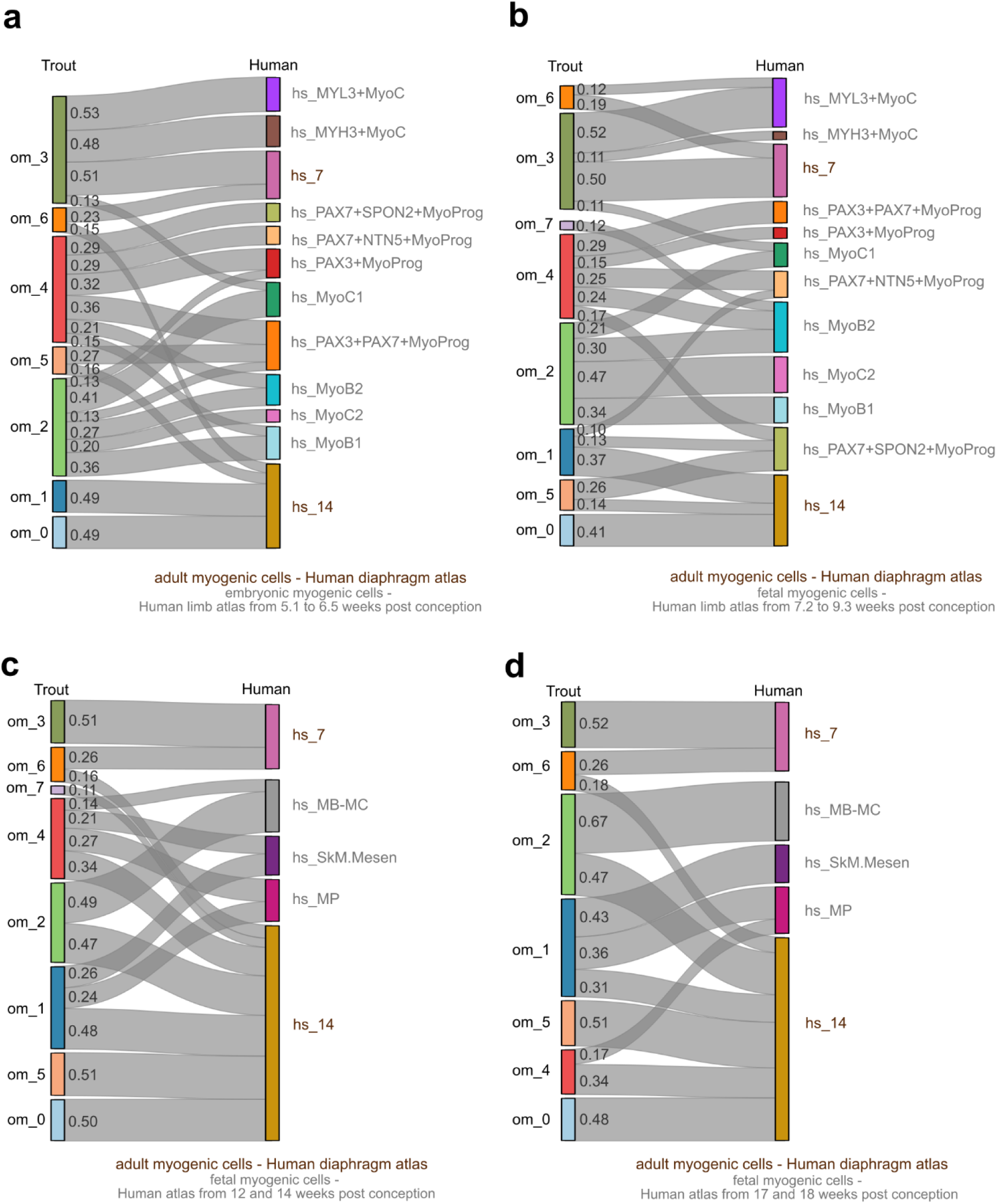
Comparative analysis of trout and human myogenic cells across developmental stages. SAMap results of myogenic cells from both species with human adult [37], embryonic [39] and fetal [38] cell atlas. Cells were colonized by cluster annotation from human (hs) or trout (om) datasets. Original annotation was kept: hs_7 and hs_14 corresponded to fast muscle cells and to skeletal muscle satellite stem cells in human adult muscle-derived cells (MDC) atlas. hs_PAX3^+^MyoProg and hs_PAX3^+^PAX7^+^MyoProg correspond to first myogenesis myogenic progenitors. hs_PAX7^+^NTN5^+^MyoProg and hs_PAX7^+^SPON2^+^MyoProg correspond to second myogenesis myogenic progenitors. hs_MyoB1, hs_MyoC1 and hs_MyoB2, hs_MyoC2 correspond to myoblasts and myocytes involved in first and second myogenesis respectively. hs_MB-MC corresponded to myoblasts and myocytes, hs_MP corresponded to myogenic progenitors and hs_SkM.Mesen corresponded to muscle progenitors with mesenchymal signature from human fetal atlas. om_0 to om_7 corresponded to the 8 clusters identified in myogenic cells from trout MDC atlas (all 9,955 cells used). (a-b) Sankey diagrams linking trout myogenic cells to human myogenic cells from adult and embryonic (a, b) cell atlases with cells from first myogenesis (a) and second myogenesis (b) based on SAMap mapping scores. (c-d) Sankey diagrams linking trout myogenic cells to human myogenic cells from adult atlas combined with fetal atlases with cells from 12 and 14 weeks post-conception (c) and 17 and 18 weeks post-conception (d) based on SAMap mapping scores. The link width is proportional to the mapping score.

### A shift from myogenic to fibro-adipogenic and extracellular matrix–associated transcriptional programs occur during the muscle growth

The origins of the transcriptomic trajectories that drive a committed cell to either fuse with a pre-existing fiber or produce a new one lay in the behavior of myogenic progenitors of clusters 0 and 1. To identify the molecular changes underlying the modifications of these transcriptomic trajectories, we performed Likelihood Ratio Tests (LRT) using DESeq2 (DESeq2-LRT, BH corrected p-value < 0.05). This approach is particularly suitable for temporal follow-ups, enabling the identification of differentially expressed genes (DEG) that share the same expression patterns in relation to trout weight. Among the 1,146 and 461 DEG identified in cluster 0 (Figure 5a) and 1 (Figure 5b) respectively, we observed enrichment of genes encoding matrix proteins, among which *col6a3* and *lama4* that were down-regulated and *col5a3a* and *lama2* that were up-regulated with weight (Figure 5c). Surprisingly, *pdgfrα*, a well-known marker gene for FAP in mammals [42], was up-regulated in cluster 0 which represented the predominantly population of myogenic precursors with body weight (Figure 5d). To our knowledge, the presence of *pax7*^+^/*pdgfrα*^+^ cells in skeletal muscle has not been described in any other adult or juvenile animal model. To validate these results, we performed dual RNAscope labeling for *pax7* and *pdgfrα* on muscle sections from 100 g trout. Despite relatively low numbers (as expected at 100 g trout), the presence of such *pax7*^+^/*pdgfrα*^+^ cells was confirmed in white muscle section (Figures 5e and S3). In addition, other fibro-adipogenic genes were found as up-regulated in these cells with the decline of hyperplasia, including *vegf*, *pdgfa*, *plin2*, and *ncam1a*. Taken together, our results showed that hyperplasia decline was accompanied by a marked change in the gene expression of matrix components, as well as an up-regulation of fibro-adipogenic genes in the myogenic progenitors.

**Figure 5.**
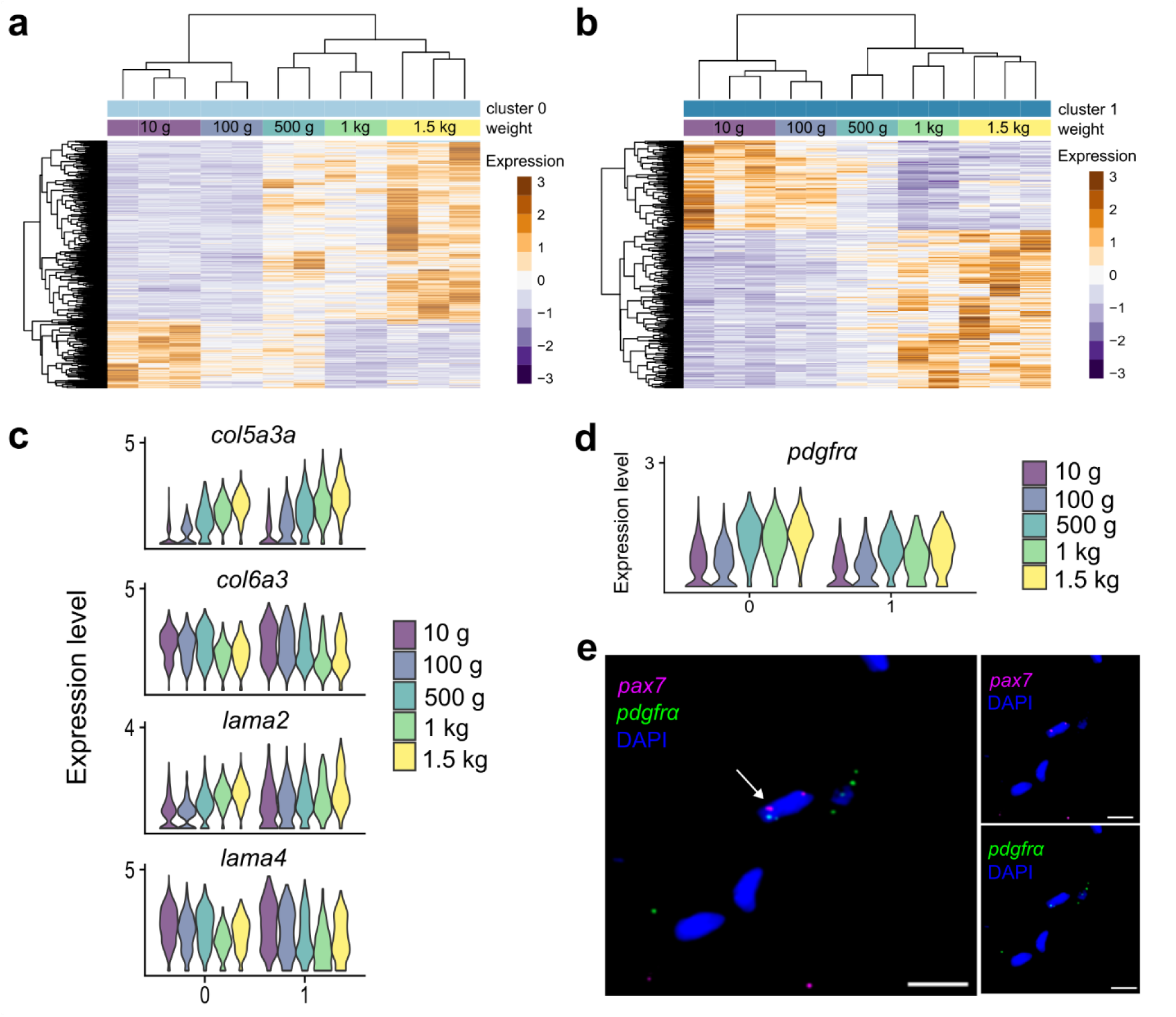
Differential expression analysis highlighted muscle stem cell plasticity and shifts of extracellular matrix composition during hyperplasia decline. (a-b) Heatmaps illustrating the differentially expressed genes in clusters 0 and 1 across weight stages. (c) Violin plot of key extracellular matrix components’ expression (including *col5a3a*, *col6a3*, *lama2* and *lama4*) in myogenic progenitors by weight. (d) Violin plot of platelet-derived growth factor receptor alpha (*pdgfrα*) expression in myogenic progenitors by weight. (c,d) The ALRA corrected matrix was used to visualize expression levels. (e) Representative images of RNAscope for *pax7* (magenta) and *pdgfrα* (green) on 100 g trout white muscle cross-sections. Arrows indicate *pax7*^+^/*pdgfrα*^+^ cell. DAPI (blue) stains the nuclei. Scale bar = 10 µm.

### Contribution of fibro-adipogenic progenitors to the muscle stem cell micro-environment shifts correlated with the decline of hyperplasia

We then investigated the second most present population in the atlas, namely mesenchymal cells that are associated with connective tissue. Our aim was to explore whether factors extrinsic to MuSC might be associated with the decline in hyperplasia. To do this, we carried out a new clustering that identified 7 clusters (Figure 6a) and submitted them to MAST test-based differential analysis (Figure 6b, Table S4). Cluster 0 was composed of chondrocytes as shown by its specific expression for *sox9* [48]. Clusters 1 and 3 were defined by a preferential expression of *pdgfrα* and proto-oncogenes as such as *jun*, suggesting that FAP [49] were found in these two clusters (Figure 6b, Table S4). The marked expression of gene coding *hes1* in cluster 3 suggested that these cells were in a state closer to quiescence. *Cathepsin-K* (*ctsk*) and *ccn4* (also known as WISP1) genes were markers of cluster 2, which shared many markers of clusters 4 and 0. This suggested that they contain differentiated cells with at least a bipotency towards chondrogenic and adipogenic lineages. Cluster 4 was characterized by its expression of the genes coding *perilipin-2* (*plin2*) and *pparg*, defining them as pre-adipocytes [50]. The specific expression of *col2a1b* and *fibronectin* (*fn1*) on the one hand and *myl2* on the other identified cells of cluster 5 and 6 as fibroblasts and myo-fibroblasts, respectively. During muscle growth, the proportion of each cluster remained equivalent except for those of cluster 2 (bipotent FAP) that decreased and cluster 3 (FAP expressing quiescent markers) and cluster 4 (pre-adipocytes) that increased (Figures 6c and S8). We performed a differential analysis (DESeq2 LRT, BH corrected p-value < 0.05) on quiescent and activated FAP (clusters 1 and 3, respectively), which are known to be major contributors to the extracellular matrix, in an attempt to find DEG according to weight and hyperplasia decline. We identified 881 and 675 DEG in cluster 1 and 3 respectively, as illustrated by the heatmaps in Figure 6d and 6e. Interestingly, 12 DEG coding for collagen proteins and 3 laminins were found, including those deregulated in myogenic precursors (Figure 6f and g). Overall, these data are consistent with a rearrangement of matrix components during muscle hyperplasia decline.

**Figure 6.**
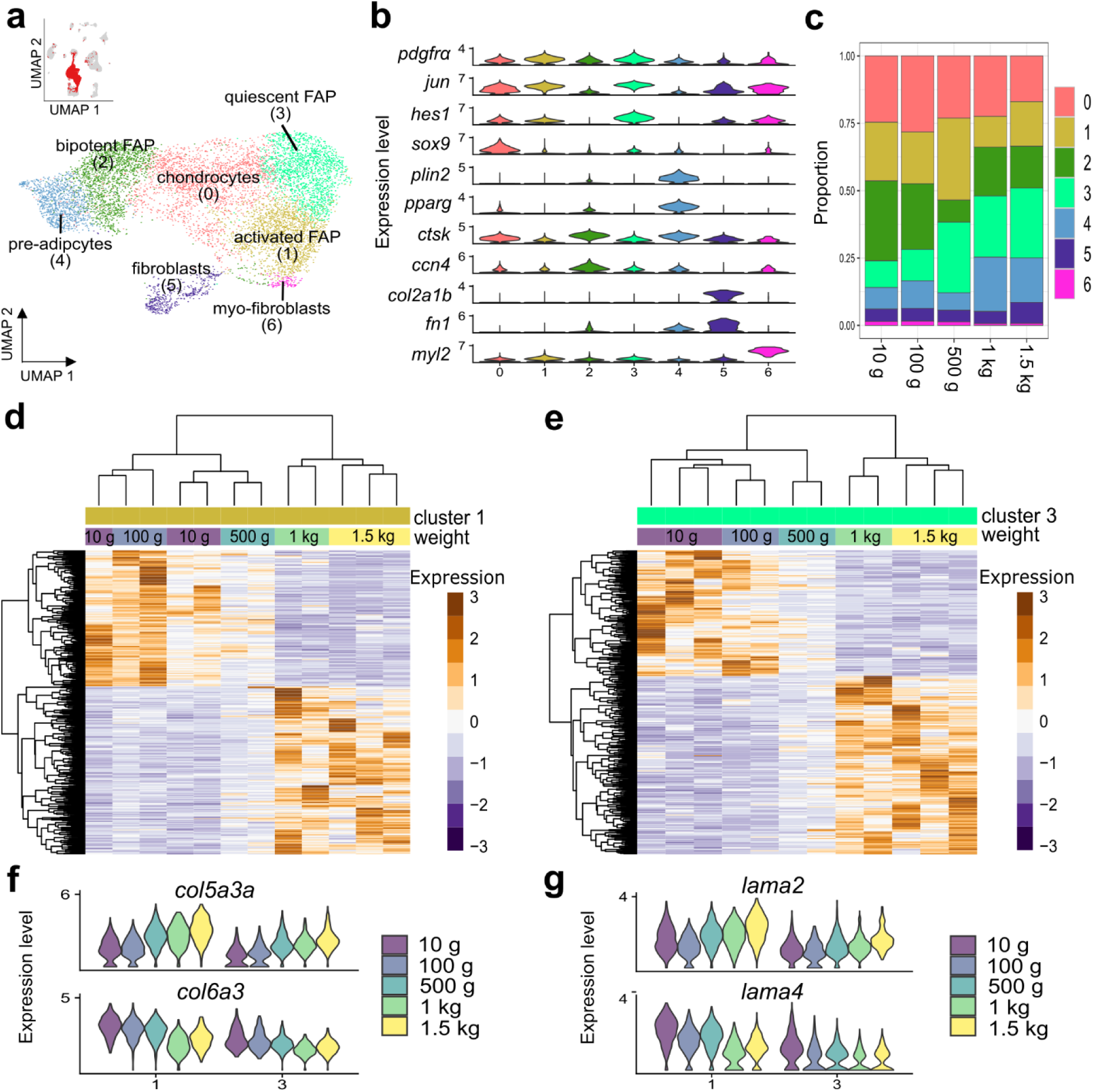
Contribution of fibro-adipogenic progenitors to changes in extracellular matrix properties. (a) UMAP of mesenchymal cells and clustering identifying 7 subpopulations (all samples and all stages merged; 9,026 cells). (b) Gene expression profiles defining mesenchymal cell subpopulations. (c) Evolution of the proportion of mesenchymal cell subpopulations during growth. (d, e) Heatmaps illustrating the differentially expressed genes in clusters 1 and 3, corresponding to quiescent and activated fibro-adipogenic progenitors (FAP) according to body weight. (f, g) Violin plot of expression of key extracellular matrix components (including *col5a3a*, *col6a3*, *lama2* and *lama4*) in primitive FAP according to body weight. (b, f, g) The ALRA corrected matrix was used to visualize expression levels.

## DISCUSSION

In mammals, muscle growth after birth results from fiber hypertrophy, whereas in most teleost such as trout, it results from both hyperplasia and hypertrophy during the juvenile period. To our knowledge, this present study provides the first single-cell resolution view of the post-hatching muscle growth at the same time as another work [51]. Our study highlights the presence of distinct myogenic transcriptional trajectories, the marked decline of a myogenic subpopulation in conjunction with the cessation of hyperplasia as a growth modality, and finally the presentation of a previously unrecorded fraction of *pax7*^+^/*pdgfrα*^+^ cells. These findings extend our understanding of MuSC diversity and its context-dependent dynamics, particularly during muscle growth with simultaneous hyperplasia and hypertrophy.

Comparing our findings to mammalian muscle single-cell atlases further underscores the distinctiveness of trout MDC diversity. In mammals, including humans, single-cell transcriptomic studies have consistently captured a low myogenic precursor proportion, particularly myoblasts, which are often underrepresented relative to non-myogenic cell types [19,52]. For example, De Micheli *et al*. (2020) identified two bifurcated states of MuSC in adult human skeletal muscle but reported limited representation of myoblasts and early differentiating progenitors [19]. Interestingly, we found that trout myoblast proportion decreases with the decline of hyperplasia, becoming closer to what is observed in adult human MDC atlas. Deeper analysis on the myogenic cell diversity revealed eight transcriptionally distinct myogenic subpopulations in trout, underscoring the complexity of cell states during trout muscle growth. Distinct myogenic cell subpopulations were found in distinct muscles [16], during regeneration [18], in myopathy [53] or during aging [54] in mammals, reminiscent of the myogenic cell heterogeneity we found. This dynamic heterogeneity is further supported by recent studies demonstrating the plasticity of MuSC to both intrinsic and extrinsic cues [21,55].

The identification of two transcriptional trajectories of myogenic cells correlated with hyperplasia and hypertrophy raises the possibility of a new regulatory paradigm. These results align with very recent findings on scRNA-seq in human limb development [39]. This study revealed multiple waves of myogenesis marked by distinct transcriptional profiles, with Pax3^+^/Pax7^+^ progenitors giving rise to spatially specialized trajectories for embryonic and fetal myogenesis. The early progenitors exhibited spatiotemporal dynamics similar to the proliferative and differentiative clusters we identified in trout, particularly in the transition from multiple intermediate phases to one late differentiating phase. In addition, myogenic subpopulations transitory present during muscle regeneration exhibit also distinct transcriptomic trajectories [18,39]. Interestingly, the authors identified a “committed” and a “cycling” cell fate, both leading to differentiated cells. These two fates are reminiscent of the transcriptional trajectories of trout myogenic cells correlated with either hyperplasia or hypertrophy. Taken together, these results show that a highly proliferative myogenic subpopulation is present only in muscles undergoing hyperplasia, *i.e*., during fetal development, regeneration, or during juvenile trout growth, which is consistent with a role in new fiber formation. Furthermore, the description of distinct transcriptional trajectories during phases of myogenesis suggests conserved regulatory frameworks in vertebrate muscle development.

Our scRNA-seq analyses and subsequent RNAscope validation reveal distinct patterns for *myod* paralogs in the myogenic program of trout, aligning with and extending prior findings in salmonids [56–58]. The four paralogs of *myod* (*myod1a*, *1b*, *2a*, and *2b*) arising from genome duplication events [59] exhibit spatially and temporally distinct expression patterns. For *myod2* paralogs, their expression in early and intermediate states of myogenic differentiation evoke more dynamic and context-specific roles. *myod2b* is preferentially expressed in intermediate myogenic cells associated with cluster 5 in our single-cell data. These cells define a trajectory that persists during hypertrophic growth. Conversely, *myod2a* is expressed in activated progenitors co-expressing myf5 (cluster 1) and proliferative myoblasts (cluster 4), associated with hyperplastic growth phase and decreasing with age. These findings agree with observations that *myod2* paralogs are tightly linked to proliferative zones and stratified hyperplasia in trout or in proliferative myoblasts in Atlantic salmon [56–58]. Interestingly, the sequential expression of *myod1* prior to *myod2* during embryogenesis in trout [56] contrasts with the adult-specific expression patterns as *myod1* was preferentially expressed in late differentiating myoblasts. We speculated that these temporal shifts reflect a divergence in regulatory pathways between embryogenesis and MuSC-mediated muscle growth. Collectively, these findings highlight the functional divergence and complementarity of *myod* paralogs, reinforcing their utility in dissecting distinct myogenic trajectories. In this sense, comparative studies in zebrafish further underscore the evolutionary plasticity of the myogenic regulatory factors or cell cycle gene network in adapting to species-specific fiber formation or growth strategies [60,61].

The transcriptional changes in ECM components during the transition from hypertrophy-hyperplasia to hypertrophy bring up the possibility of niche remodeling in regulating MuSC behavior during growth. Very recently, a study identified several ECM components that were differentially expressed by MuSC in different species undergoing distinct levels of hyperplasia [51]. Of note, MuSC-specific depletion of the *col4a2* gene was shown to alter their dynamics in zebrafish [51]. In the present study, we found that both MuSC and FAP presented a shift in the matrix component expression in favor of an autonomous [62] and non-autonomous [63] effect on MuSC behavior. It is also possible that the shift in FAP transcriptional signature could be a reflect of an increased requirement for stronger tissue connectivity in larger fish, which generate higher overall forces during locomotion. Although ECM proteins such as laminins and collagens have been shown to influence MuSC fate and myogenic commitment in mammals [64–67], our study highlights a correlation, while causality remains to be further explored. A population of FAP has been shown to express *ccn4* (also known as WISP1). It is interesting to note that this factor, promoting myoblast proliferation [68], is a marker of bipotent FAP that fall as hyperplasia declines. In mammals, the dynamic interplay between MuSC and their niche is further supported by growth factors and cytokines, including VEGF and PDGF [23]. We observed an increase of these factors during hyperplasia decline, suggesting that similar mechanisms operate in trout, facilitating the transition between growth modalities. We demonstrated *in situ* the existence of *pdgfrβ*^+^ and *pax7*^+^/*pdgfrβ*^+^ cells associated with the hypertrophic growth trajectory. It will be very interesting to investigate if these cells are localized near capillaries regarding the increase of endothelial cell proportions we determined with weight. Another notable result is the up-regulation of *pdgfrα* indicating a shift in MuSC plasticity that may promote fibro-adipogenic commitment as hyperplasia declines. Expression of *pdgfrα*, along with other mesenchymal markers, in myogenic progenitor subset has been previously reported in embryonic and fetal human muscle [38]. In post-natal mammalian muscle, such cells have also recently been described, but only at the myotendinous junction [69, 70]. This contrasts with our data showing these cells sparsely distributed throughout the tissue, and not predominantly at the myoseptal junction. To our knowledge, only two other studies mention the presence of these cells that correspond to bipotent progenitors capable of contributing to both myogenic and connective tissue lineages during embryonic development [71] and in aging [72]. In the second case, a notion of conversion of MuSC from myogenic to fibrogenic lineage in older animals has been put forward. This reinforces the idea that similar mechanisms may regulate MuSC plasticity and niche remodeling across vertebrates.

Recent studies in mammalian models have highlighted the interplay between MuSC and immune cells in shaping the regenerative niche [24, 73]. Exploring similar interactions in trout could provide new insights into the cross-talk between myogenic and non-myogenic cells during muscle growth. Among non-myogenic cells, we found an increased proportion of endothelial cells over time. It could be informative to focus on cross-talk existing between them and MuSC during trout muscle growth, especially as these cells have been described as potential close partners providing systemic factors and cytokines [74]. Future works will be required to produce a complementary single nuclei RNA sequencing analysis to provide a complete story about MuSC and their niche in the decline of hyperplastic growth. Indeed, at this stage, the muscle fiber, despite being a major player in the stem cell micro-environment [75], is not taken into account when deciphering the growth process, as it is absent from scRNA-seq. From an evolutionary perspective, the trout model offers a unique opportunity to study the maintenance of hyperplastic growth in teleost. Comparative analyses with other teleost species that exhibit different growth strategies during development and after hatching could enhance our understanding of muscle formation in vertebrates [8, 76].

## LIMITATIONS OF THE STUDY

In the present study, several limitations should be considered. First, our atlas relies on droplet-based scRNA-seq of dissociated cells isolated from fast (white) muscle; therefore, it does not capture multinucleated myofibers and may miss progenitor and precursor populations located outside this muscle region. Second, because mature fibers are absent from scRNA-seq, myofiber-derived cues were inferred indirectly; moreover, the “fast myocyte” cluster may include a fraction of inadvertently captured myonuclei released during the tissue dissociation step. Considering that cell suspensions were stored overnight at 4°C prior to single cell processing, it cannot dismiss the idea that stress-related transcriptional changes might be induced despite identical handling across conditions. Our study is cross-sectional across weight stages. Indeed, as body weight covaries with other physiological parameters (*i.e*., age, endocrine status and mechanical load), the relationships we report between cell-state dynamics, niche remodeling and growth modalities remain correlative and will require functional validation. Finally, cross-species SAMap alignments depend on orthology/paralogy assignments and should be interpreted as homology-guided correspondences rather than definitive equivalences. Future work combining functional perturbations, spatial approaches and complementary snRNA-seq, as well as comparative analyses across teleosts with distinct growth strategies, will be required to refine the mechanisms controlling postnatal muscle growth.

## CONCLUSIONS

Overall, our results raise the possibility of the existence of two myogenic trajectories with distinct dynamics correlated with post-hatching muscle hyperplasia and hypertrophy. The discovery of this two-branch myogenic trajectory and a tissue-resident *pax7*^+^/*pdgfrα*^+^ cell population in trout provides useful framework for investigating the regulatory mechanisms of muscle growth and regeneration across vertebrates. These findings raise several questions about the functional roles of these cells in different physiological contexts, including responses to injury and environmental stressors. It will be essential to investigate the signaling pathways that regulate their dynamics and contributions to muscle growth under different conditions. Additionally, integrating single-cell transcriptomics with epigenomics could uncover regulatory networks driving cell fate decisions, as demonstrated in recent studies on muscle regeneration or aging [29, 77, 78].

## Supporting information

Figure S1: RNAscope negative controls.

Figure S2: Representative images of RNAscope for pax7 (magenta), myod1 (green) or myod2a (white) on 100 g trout white muscle cross-sections.

Figure S4: Homology of muscle-derived cells between trout and human atlases.

Figure S5: Genes associated with myogenic cell fates in juvenile trout.

Figure S6: Weight-dependent changes in myogenic cells subset composition from muscle-derived cell trout atlas.

Figure S7: Cross-species SAMap integration of juvenile trout myogenic cells with combined adult and fetal human myogenic cells from skeletal muscle at

Figure S8: Weight-dependent changes in composition of mesenchymal cell subset from muscle-derived cell trout atlas.

Table S1: Table of sample metadata.

Table S2: Table of Official NCBI Gene Names, Gene Description and gene names used in this article.

Table S3: Table of top 100 of gene makers of myogenic cells clustering.

Table S4: Table of top 100 of gene makers of mesenchymal cells clustering.

Table S5: Mapping scores from SAMap analysis mononuclear cell from trout muscle vs. mononuclear cell from adult human muscles.

Table S6: Table of mapping scores from SAMap analysis of myogenic cells from trout muscles vs. adult and 5.1 to 6.5-week-old embryonic human muscles.

Table S7: Table of mapping scores from SAMap analysis of myogenic cells from trout muscles vs. adult and 7.2 to 9.3-week-old embryonic/early fetal hum

Table S8: Table of mapping scores from SAMap analysis of myogenic cells from trout muscles vs. adult and 12/14-week-old fetal human muscles.

Table S9: Table of mapping scores from SAMap analysis of myogenic cells from trout muscles vs. adult and 17/18-week-old fetal human muscles.

Table S10: Table of gene drivers from UniTVelo modelling classify in induction, repression or transient states.

## ABBREVIATIONS

MDC: Muscle Derived Cells
RNA: Ribonucleic Acid
MuSC: Muscle Stem Cells
SC: Satellite Cells
scRNA-seq: Single-Cell RNA sequencing
FAP: Fibro-Adipogenic Progenitors
UMAP: Uniform Manifold Approximation and Projection
SAMap: Self-Assembling Manifold mapping
LRT: Likelihood Ratio Tests
DEG: Differentially Expressed Genes
ECM: Extra-Cellular Matrix
cDNA: complementary Deoxyribonucleic Acid
UMI: Unique Molecular Identifiers
FFPE: Formalin-Fixed Paraffin-Embedded

## DECLARATIONS

### Ethics approval and consent to participate

Rainbow trout (Oncorhynchus mykiss) were reared at the INRAE Fish Physiology and Genomic Laboratory (LPGP) experimental facilities (https://doi.org/10.15454/45d2-bn67, permit number D35-238-6, Rennes, France) under natural simulated photoperiod, at 12 ± 1°C (pH 7.8–8.4; NH4 < 0.1 mg/L) and fed ad libitum (commercial diet). Fish were anesthetized and euthanized with tricaine (MS-222) at 50 mg/L and 200 mg/L, respectively. All experimental procedures were carried out in strict accordance with the European Directive 2010/63/EU on the protection of animals used for scientific purposes. The euthanasia procedure was approved by the Ethical Committee for Animal Experimentation of Rennes (CREEA) and received the approval of French minister of national education, research and innovation under the authorization number: APAFIS #2015121511031837.

### Authors’ contributions

S.J, C.B, K.R and J.C.G conceptualized the study; S.J and N.S performed cell extraction; S.J and C.B realized single-cell RNA-seq experiments; C.B performed RNAscope experiments; S.J carried out single-cell RNA-seq analysis; S.J, C.B, K.R and J.C.G interpreted the data; S.J, C.B, K.R and J.C.G acquired funding; S.J, C.B, F.L.G, K.R and J.C.G drafted and critically reviewed the manuscript. All authors have read and approved the final manuscript.

## Acknowledgements

We thank the LPGP fish facilities (https://doi.org/10.15454/45d2-bn67) and particularly C. Duret for trout rearing and egg production. The authors acknowledge the Cytocell - Flow Cytometry and FACS core facility (SFR Bonamy, BioCore, Inserm UMS 016, CNRS UAR 3556, Nantes, France) for its technical expertise and help, member of the Scientific Interest Group (GIS) Biogenouest and the Labex IGO program supported by the French National Research Agency (n°ANR-11-LABX-0016-01). We are most grateful to the Genomics Core Facility GenoA, member of Biogenouest and France Genomique and to the Bioinformatics Core Facility BiRD, member of Biogenouest and Institut Français de Bioinformatique (IFB) for the use of their resources and their technical support. We thank Pascal Maire (Université de Paris, Institut Cochin, INSERM, CNRS, Paris, France) and Olivier Pourquié (Department of Pathology, Brigham and Women’s Hospital, Department of Genetics, Harvard Stem Cell Institute, Harvard Medical School, Harvard University, Boston, USA) for helpful discussion and improving the manuscript.

## ADDITIONAL FILES

**Figure S1: RNAscope negative controls.** RNAscope stains in white muscle cross-sections of 100 g trout. Representative images showing the expression of negative DapB C1 (magenta), DapB C2 (green) and DapB C3 (white). DAPI stains the nuclei. Scale bar = 20 µm.

**Figure S2: Representative images of RNAscope for pax7 (magenta), myod1 (green) or myod2a (white) on 100 g trout white muscle cross-sections.** DAPI (blue) stains the nuclei. Scale bar = 5 µm.

**Figure S3: Representative images of RNAscope for pax7 (magenta) and pdgfrα or pdgfrβ (green) on 100 g trout white muscle cross-sections.** DAPI (blue) stains the nuclei. Scale bar = 5 µm.

**Figure S4: Homology of muscle-derived cells between trout and human atlases.** (a) UMAP showing the cross-species comparison between atlas of trout (om) and human [37] (hs) muscle-derived cells (MDC), using SAMap [40]. Cells were colorized according to original cluster annotations. (b) UMAP visualization from cross-species comparison using SAMap. Colorized cells were from trout MDC atlas. (c) UMAP visualization from cross-species comparison using SAMap. Colorized cells were from human MDC atlas. (d) Sankey plot showing the proportion of trout MDC classified according to their most similar human equivalent based on SAMap cell-type homology analysis. (e) Stacked bar plot showing the relative proportions of the eight identified cell clusters across individual biological replicates from five weight groups (10 g, 100 g, 500 g, 1 kg and 1.5 kg). Each bar represents one biological replicate, and colors correspond to annotated clusters, illustrating sample-level variability and shifts in cell-type composition with increasing body weight. (f-h) Representative boxplots showing the proportion of selected clusters across weight groups. Cluster proportions were analyzed using the scanpro framework [35]. Statistical differences across weight groups were assessed using a linear model analogous to ANOVA. P-values are indicated for each cluster (Benjamini-Hochberg adjusted). Colored frames and titles correspond to cluster identity as in panel (e).

**Figure S5: Genes associated with myogenic cell fates in juvenile trout**. (a-c) Heatmaps of predicted induction (a), repression (c) and transient (e) gene expressions are resolved along the inferred cell time, showing a clear separation in temporal space. Induction genes tend to be active at the end of the cellular process, while repression genes display the opposite behavior. (b,d) Phase portraits depicting the expression of *myod1* (b), *myod2a* (d), and platelet-derived growth factor receptor beta (*pdgfrβ*) (f), correlating gene expression (following the amount of spliced and unspliced transcripts) with latent time.

**Figure S6: Weight-dependent changes in myogenic cells subset composition from muscle-derived cell trout atlas**. (a) Stacked bar plot showing the relative proportions of the eight identified cell clusters across individual biological replicates from five weight groups (10 g, 100 g, 500 g, 1 kg and 1.5 kg). Each bar represents one biological replicate, and colors correspond to clusters (0-7), illustrating sample-level variability and shifts in cell-type composition with increasing body weight. (b-e) Representative boxplots showing the proportion of selected clusters across weight groups. Cluster proportions were analyzed using the scanpro framework [35], which models cell-type proportions at the sample level to account for biological replication. Statistical differences across weight groups were assessed using a linear model analogous to ANOVA. P-values are indicated for each cluster (Benjamini-Hochberg adjusted). Colored frames and titles correspond to cluster identity as in panel (a).

**Figure S7: Cross-species SAMap integration of juvenile trout myogenic cells with combined adult and fetal human myogenic cells from skeletal muscle atlases.** (a) UMAP visualization of adult human skeletal muscle-associated cell populations reproduced from the Tabula Sapiens atlas [37], shown to provide a reference map of adult human cell states used for cross-species comparison. Cell identities correspond to annotated populations in the original atlas. (b-c) Force-directed graph layouts of embryonic human limb myogenic cells reproduced from [39] to visualized cell types (b) and ages in weeks post-conception (c). (d-e) UMAP embeddings generated from SAMap cross-species integration between trout juvenile myogenic cells and myogenic cells from human embryonic and adult datasets with cells from first myogenesis (d) and second myogenesis (e). Human cell annotations are preserved from the original studies, allowing direct visual comparison with the published reference atlases. (f-g) t-SNE representations of human fetal skeletal muscle cells at later fetal stages reproduced from [38]. (f) 12-14 weeks post-conception. (g) 17-18 weeks post-conception. These plots provide developmental reference structure for late fetal myogenic states. (h-i) UMAP embeddings generated from SAMap cross-species integration between trout juvenile myogenic cells and myogenic cells from human fetal stages correspond to 12-14 weeks post-conception (h) and 17-18 weeks post-conception (i), and adult datasets displaying integrated trout and human cell populations colored by original cluster identity.

**Figure S8: Weight-dependent changes in composition of mesenchymal cell subset from muscle-derived cell trout atlas**. (a) Stacked bar plot showing the relative proportions of the eight identified cell clusters across individual biological replicates from five weight groups (10 g, 100 g, 500 g, 1 kg and 1.5 kg). Each bar represents one biological replicate, and colors correspond to clusters (0-6), illustrating sample-level variability and shifts in cell-type composition with increasing body weight. (b-d) Representative boxplots showing the proportion of selected clusters across weight groups. Cluster proportions were analyzed using the scanpro framework [35], which models cell-type proportions at the sample level to account for biological replication. Statistical differences across weight groups were assessed using a linear model analogous to ANOVA. P-values are indicated for each cluster (Benjamini-Hochberg adjusted). Colored frames and titles correspond to cluster identity as in panel (a).

**Table S1: Table of sample metadata.**

**Table S2: Table of Official NCBI Gene Names, Gene Description and gene names used in this article.**

**Table S3: Table of top 100 of gene makers of myogenic cells clustering.**

**Table S4: Table of top 100 of gene makers of mesenchymal cells clustering.**

**Table S5: Mapping scores from SAMap analysis mononuclear cell from trout muscle vs. mononuclear cell from adult human muscles.** Mapping table with a legend table: cluster number from trout (om) or human (hs) and their cell type annotation.

**Table S6: Table of mapping scores from SAMap analysis of myogenic cells from trout muscles vs. adult and 5.1 to 6.5-week-old embryonic human muscles.**

**Table S7: Table of mapping scores from SAMap analysis of myogenic cells from trout muscles vs. adult and 7.2 to 9.3-week-old embryonic/early fetal human muscles.**

**Table S8: Table of mapping scores from SAMap analysis of myogenic cells from trout muscles vs. adult and 12/14-week-old fetal human muscles.**

**Table S9: Table of mapping scores from SAMap analysis of myogenic cells from trout muscles vs. adult and 17/18-week-old fetal human muscles.**

**Table S10: Table of gene drivers from UniTVelo modelling classify in induction, repression or transient states.**

