## Supplementary figures and images for "Distinct muscle stem cell fates correlated with hyperplasia and hypertrophy during skeletal muscle growth in rainbow trout"

### Figure S1: RNAscope negative controls.

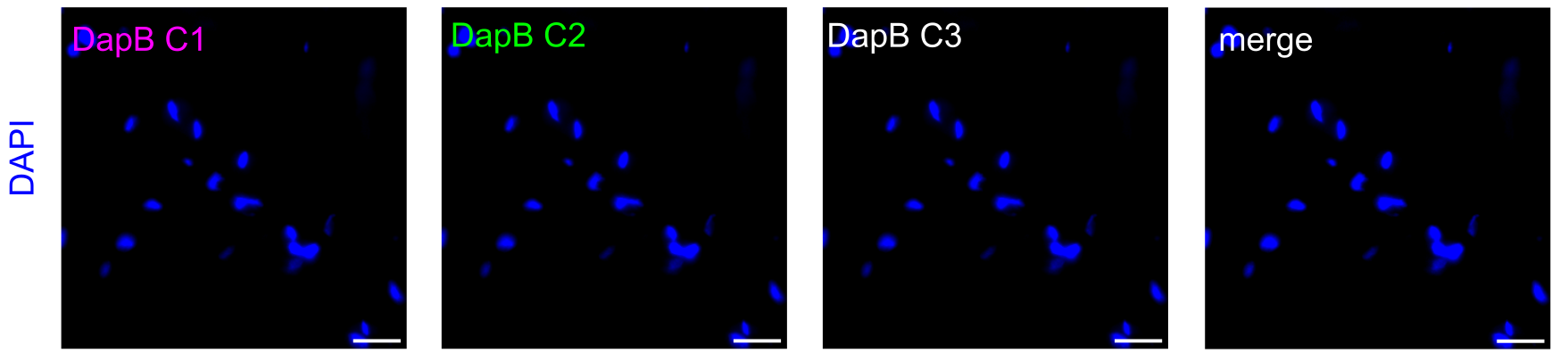

### Figure S2: Representative images of RNAscope for pax7 (magenta), myod1 (green) or myod2a (white) on 100 g trout white muscle cross-sections.

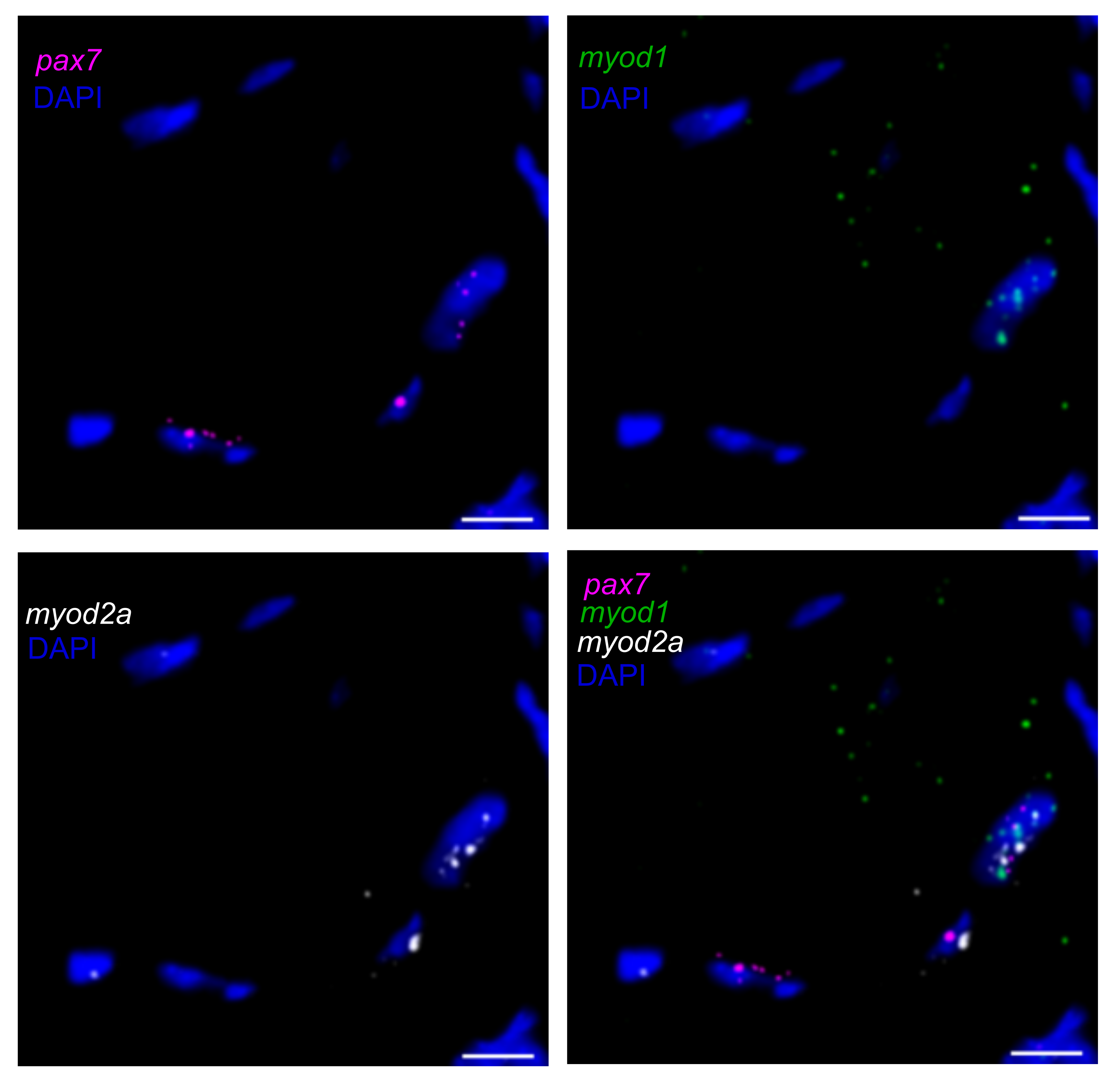

### Figure S4: Homology of muscle-derived cells between trout and human atlases.

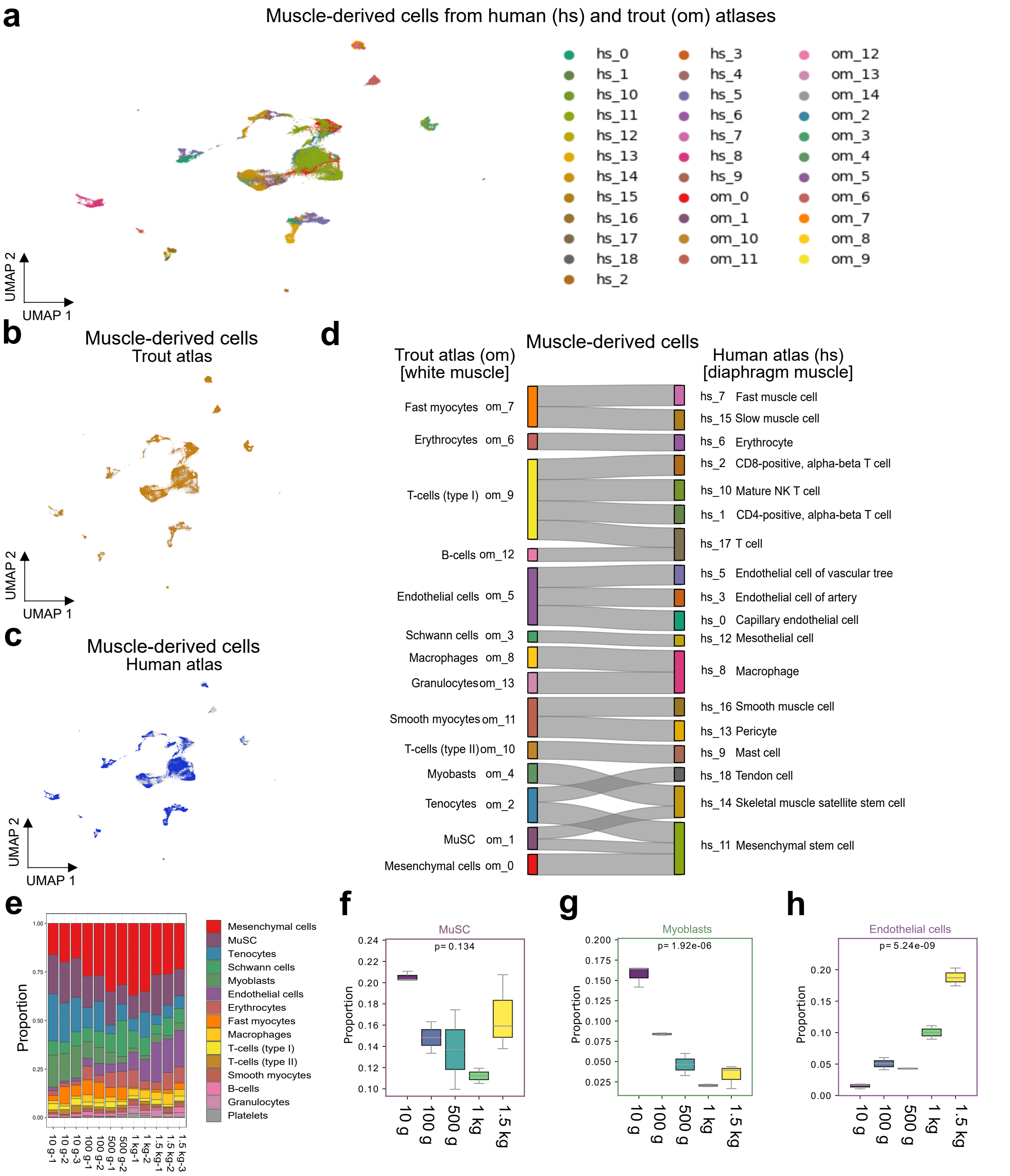

### Figure S5: Genes associated with myogenic cell fates in juvenile trout.

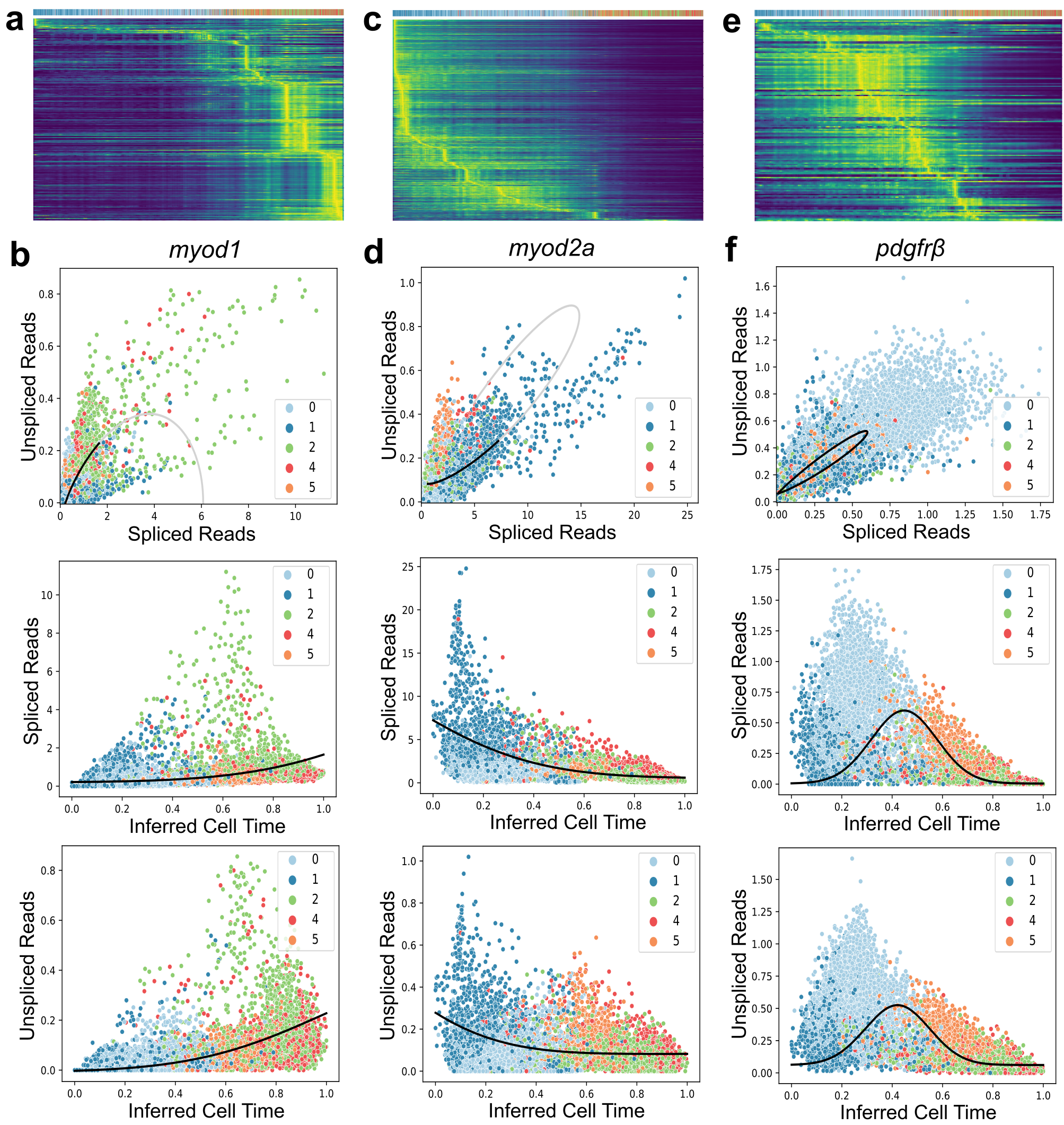

### Figure S6: Weight-dependent changes in myogenic cells subset composition from muscle-derived cell trout atlas.

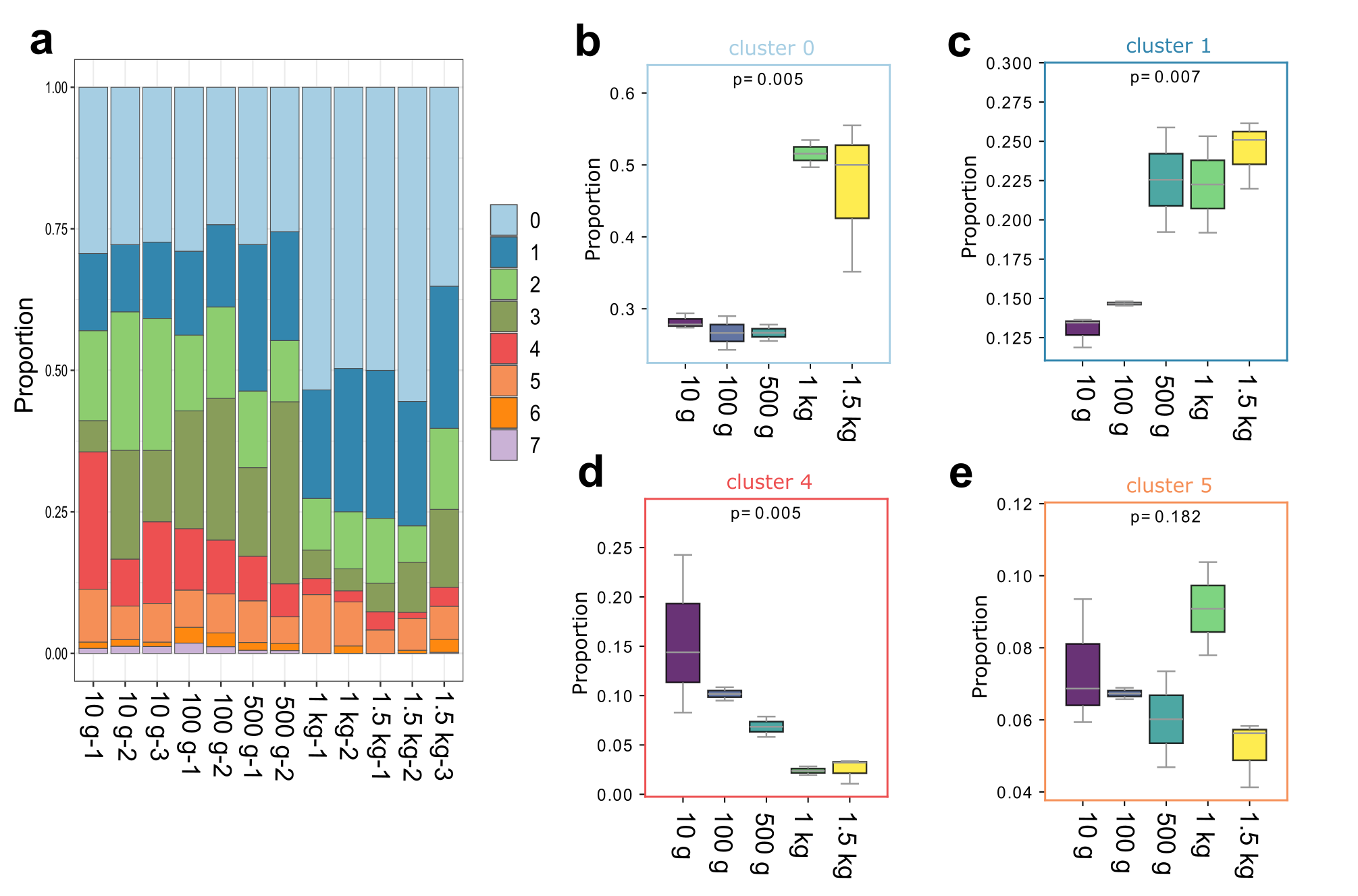

### Figure S7: Cross-species SAMap integration of juvenile trout myogenic cells with combined adult and fetal human myogenic cells from skeletal muscle at

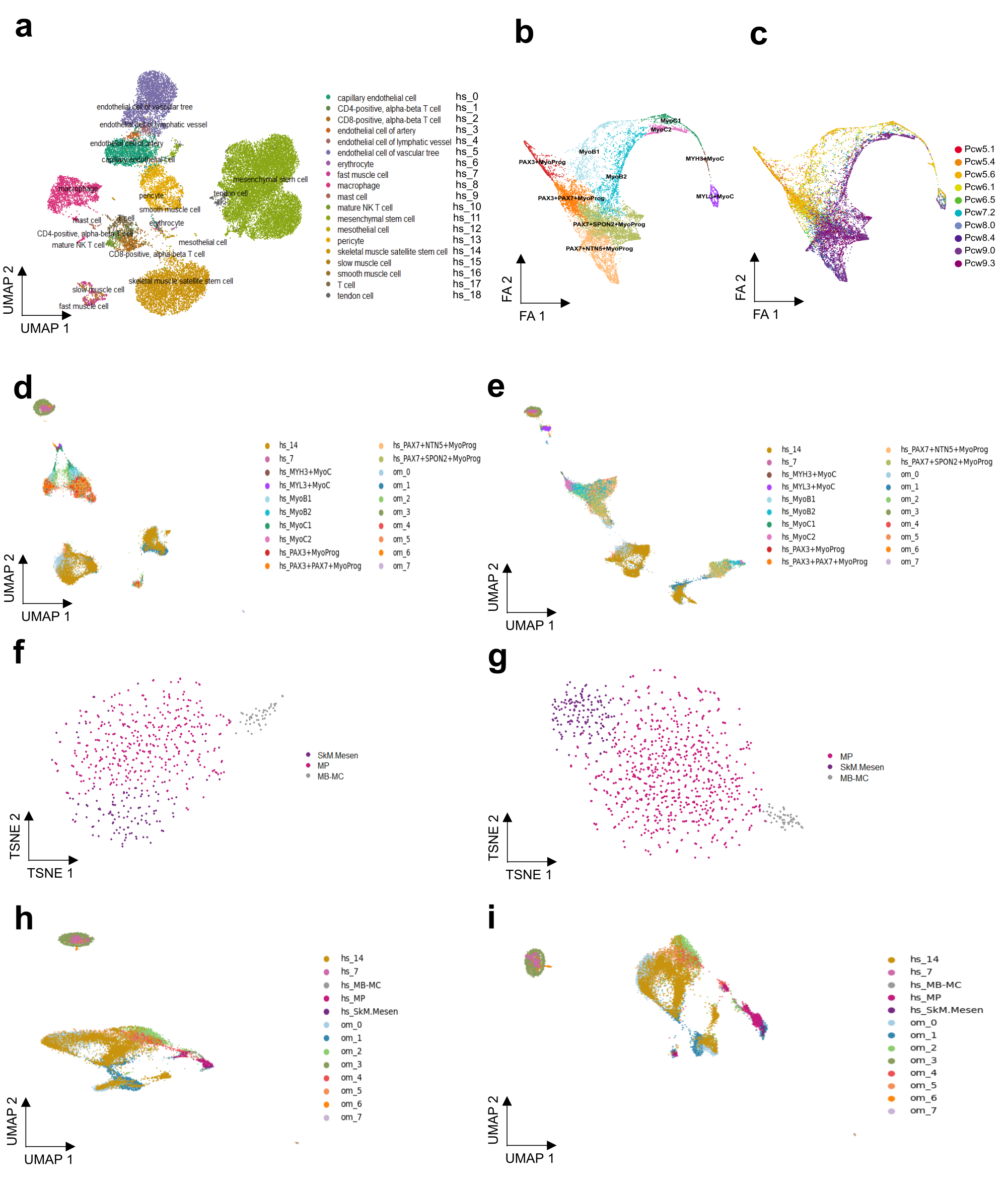

### Figure S8: Weight-dependent changes in composition of mesenchymal cell subset from muscle-derived cell trout atlas.

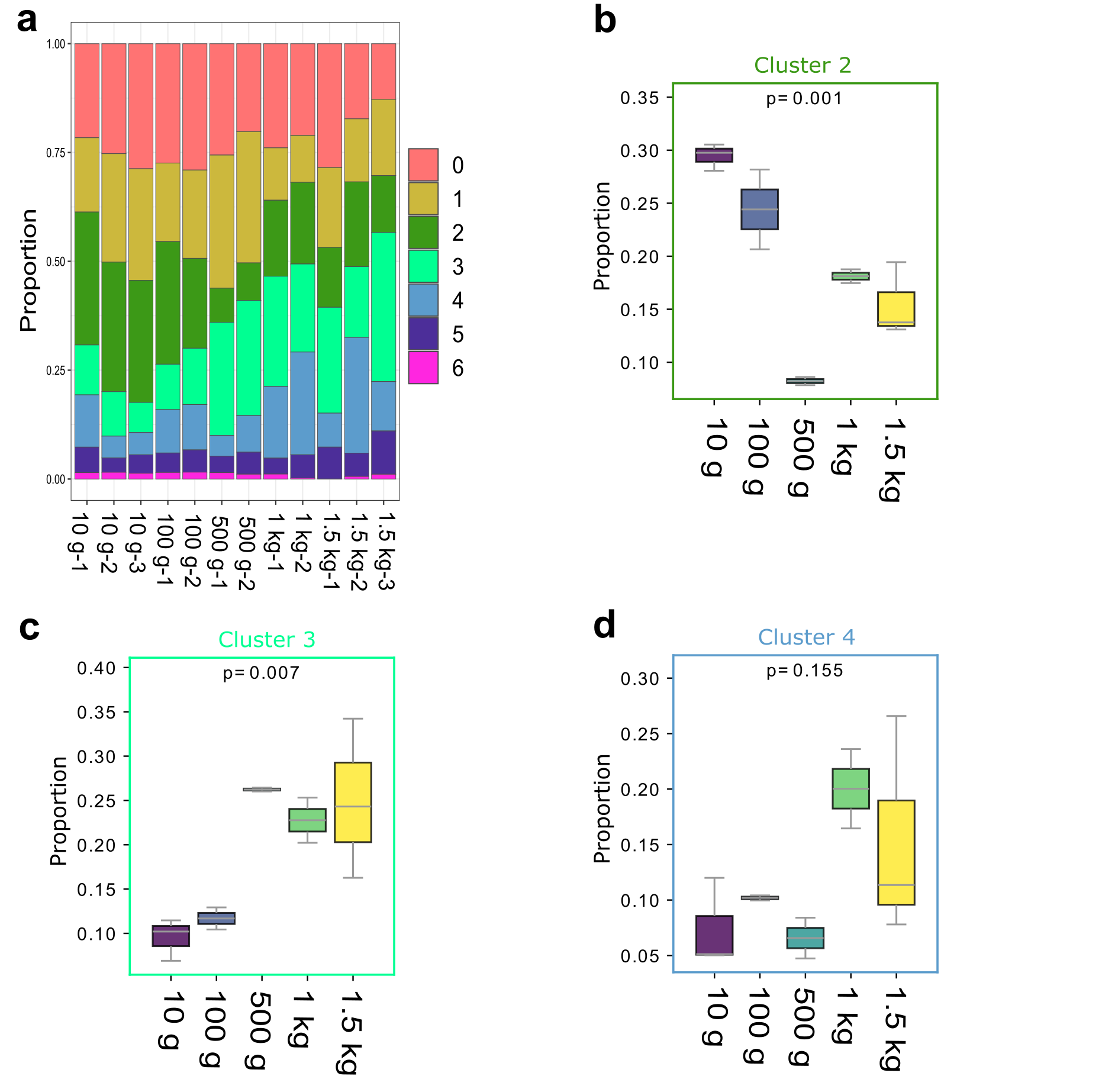

### Supplemental Data 1

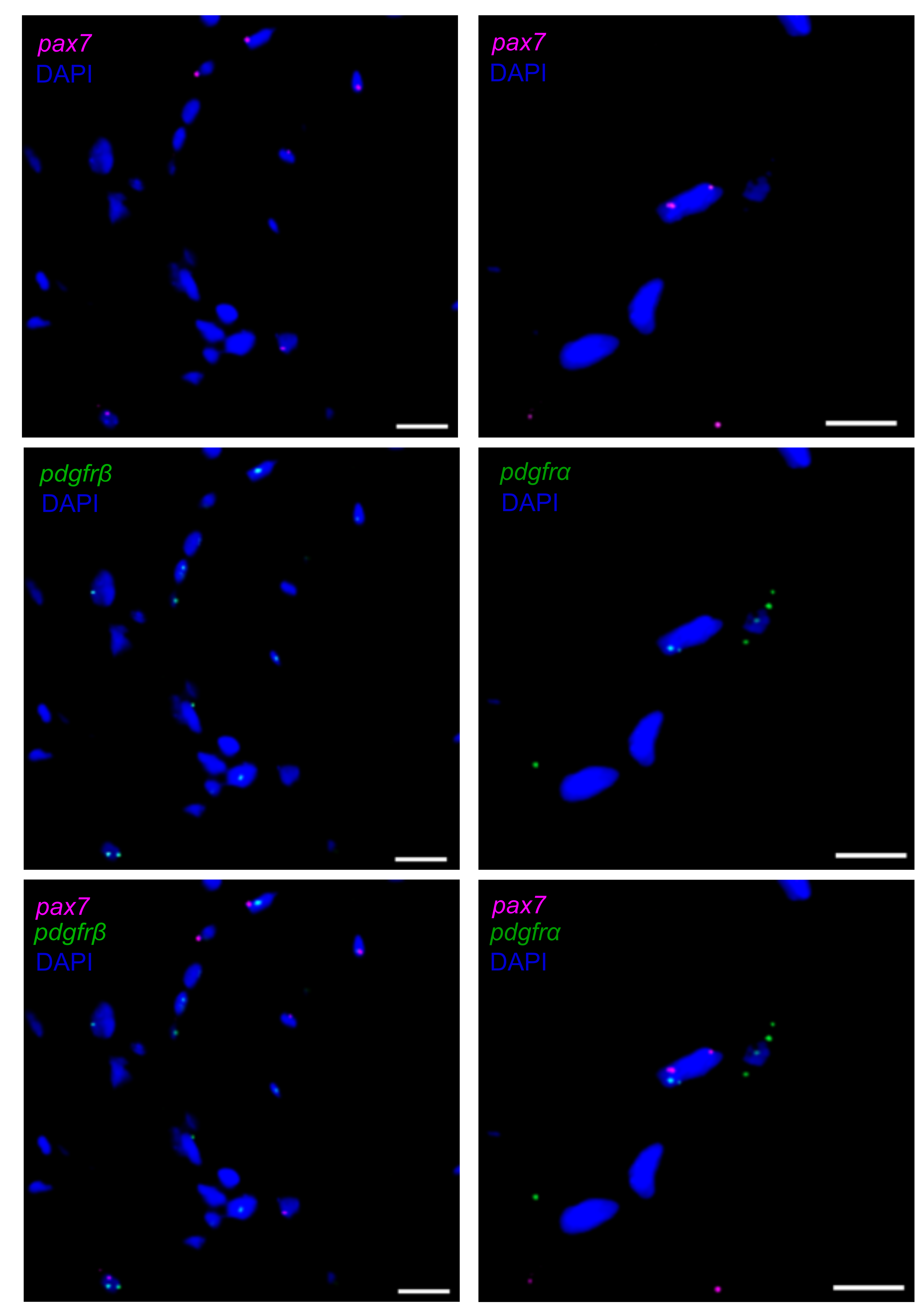
