## Supplementary material for "Distinct muscle stem cell fates correlated with hyperplasia and hypertrophy during skeletal muscle growth in rainbow trout": Table S2: Table of Official NCBI Gene Names, Gene Description and gene names used in this article.

| **Gene Name** | **Gene Description** | **Gene Name in the article** |
| --- | --- | --- |
| LOC110521062 | fibronectin type III domain-containing protein 1 | *fndc1* |
| pdgfra | platelet-derived growth factor receptor, alpha polypeptide | *pdgfrα* |
| LOC110528585 | integrin alpha-7 | *itga7* |
| LOC110532152 | paired box protein Pax-7 | *pax7b* |
| tnmd | tenomodulin | *tnmd* |
| mbpa | myelin basic protein a | *mbpa* |
| myog | myogenin | *myog* |
| LOC110506352 | cadherin-5 | *cdh5* |
| vwf | von Willebrand factor | *vwf* |
| hba4 | Hemoglobin subunit alpha-4 | *hba4* |
| LOC110501785 | myosin light chain 1, skeletal muscle isoform | *myl1b* |
| LOC110492629 | troponin C, skeletal muscle | *tnnc2* |
| ptprc | protein tyrosine phosphatase receptor type C | *ptprc* |
| LOC110501689 | macrophage receptor MARCO | *marco* |
| LOC118964336 | T-cell-specific surface glycoprotein CD28-like | *cd28* |
| LOC110492331 | interleukin-17 receptor B | *il17rb* |
| mustn1a | musculoskeletal, embryonic nuclear protein 1a | *mustn1a* |
| LOC110498989 | actin, aortic smooth muscle | *acta2* |
| pax-5 | Pax-5 protein | *pax5* |
| LOC100136240 | neutrophil cytosolic factor 2 | *ncf2* |
| LOC110525802 | platelet glycoprotein Ib beta chain | *gp1bb* |
| LOC110490466 | myogenic factor 5-like | *myf5* |
| mki67 | marker of proliferation Ki-67 | *mki67* |
| myod2 | myoblast determination protein 2 | *myod2a* |
| LOC110493125 | myoblast determination protein 1 homolog 2-like | *myod2b* |
| myod | MYOD protein | *myod1* |
| mymk | myomaker, myoblast fusion factor | *mymk* |
| myl1 | myosin, light chain 1, alkali; skeletal, fast | *myl1a* |
| LOC110521616 | Jun proto-oncogene, AP-1 transcription factor subunit | *jun* |
| LOC110534079 | transcription factor HES-1 | *hes1* |
| sox9 | SOX9 alpha2 | *sox9* |
| LOC110500648 | perilipin-2 | *plin2* |
| pparg | peroxisome proliferator-activated receptor gamma | *pparg* |
| LOC110514697 | cathepsin K | *ctsk* |
| LOC110496658 | CCN family member 4 | *ccn4* |
| col2a1b | collagen, type II, alpha 1b | *col2a1b* |
| LOC110528732 | fibronectin | *fn1* |
| LOC100136718 | fast myosin light chain 2 | *myl2* |
| pax7a | paired box 7a | *pax7a2* |
| LOC110530949 | paired box protein Pax-3-like | *pax3-like* |
| LOC110538970 | collagen, type V, alpha 3a | *col5a3a* |
| LOC110504831 | platelet-derived growth factor receptor beta | *pdgfrβ* |
| LOC110501275 | collagen alpha-3(VI) chain | *col6a3* |
| LOC100136119 | platelet-derived growth factor subunit A | *pdgfa* |
